# The Molecular Basis of Differentiation Wave Activity in Embryogenesis

**DOI:** 10.1101/2024.06.04.597397

**Authors:** Bradly Alicea, Surosh Bastani, Natalie K. Gordon, Susan Crawford-Young, Richard Gordon

## Abstract

As development varies greatly across the tree of life, it may seem difficult to suggest a model that proposes a single mechanism for understanding collective cell behaviors and the coordination of tissue formation. Here we propose a mechanism called differentiation waves, which unify many disparate results involving developmental systems from across the tree of life. We demonstrate how a relatively simple model of differentiation proceeds not from function-related molecular mechanisms, but from so-called differentiation waves. A phenotypic model of differentiation waves is introduced, and its relation to molecular mechanisms is proposed. These waves contribute to a differentiation tree, which is an alternate way of viewing cell lineage and local action of the molecular factors. We construct a model of differentiation wave-related molecular mechanisms (genome, epigenome, and proteome) based on *C. elegans* bioinformatic data. To validate this approach across different modes of development, we evaluate protein expression across different types of development by comparing the nematode *Caenorhabditis elegans* with several model organisms: fruit flies (*Drosophila melanogaster*), yeast (*Saccharomyces cerevisiae*), and mouse (*Mus musculus*). Inspired by gene regulatory networks, two Models of Interactive Contributions (fully-connected MICs and ordered MICs) are used to suggest potential genomic contributions to differentiation wave-related proteins. This, in turn, provides a framework for understanding differentiation and development.

## 1 Introduction

Understanding the process of differentiation during embryogenesis requires us to understand more than collections of genetic mechanisms. White genetic mechanisms can drive single cell differentiation and reaction-diffusion processes in tissues, we must also connect these to the mechanics and collective dynamics of tissue-level changes. Cross-species commonalities amongst various tissue types and the subsequent development of stem cells (Oppenheimer, 1939; Le Lièvre and Le Douarin, 1975; Le Douarin, 1988; Dieterlen-Lièvre and Le Douarin, 1991; Le Douarin et al., 1993; Le Douarin et al., 1994; Le Douarin et al., 1998; Teillet et al., 1999; Gordon, 2011) suggests a universal differentiation mechanism across multicellular Eukaryotes that connects the molecular action and phenotype. This requires a theoretical perspective that unifies a spatiotemporal model of embryogenesis that is grounded in molecular mechanisms. In this paper, we will pose differentiation as the product of differentiation waves, involving a range of processes ranging from cellular mechanics to molecular mechanisms. Our study will use secondary data to provide the basic contours of the differentiation by wave action: gene expression and protein abundance becomes relevant to expansion and contraction events in pre-differentiated cell populations.

The process of regional differentiation (Slack, 1991; Brugmann et.al, 2007; Micali et.al, 2023) is spatially explicit, while also being lineage tree and species-specific. While a few of these regulatory factors have large effect sizes (MyoD/Myf5 for muscle cell differentiation in Mammals, see Rudnicki et.al, 1993; Pou5/Oct4 control of pluripotency in Mammals, see Sukparangsi et.al, 2022), many of them have small effect sizes. Aside from the typical function-specific genes, regional differentiation involves proteins relevant to the developmental process such as Notch (Micali et.al, 2023) and Wnt (Brugmann et.al, 2007). Anterior-posterior polarity in both *C. elegans* and *Drosophila* (Kimelman and Martin, 2012), along with more specific organizational changes in the cellular architecture such as the *C. elegans* germ line (P4 sublineage, see Seidel et.al, 2018) require a process that identify collective behavior amongst cells in a regionally-specific manner. From a temporal perspective, regional differentiation processes provide new functional and structural components that punctuate the dynamics of embryonic gene expression. Single-cell approaches may not capture the complexity needed here. For example, cell atlases focus on using candidate genes for specific cell types to bolster support for cell lineage trees (Domcke and Shendure, 2023). But they do not help us identify the transformative processes themselves, which involve mechanisms such as synchronization of cell populations, the remodeling of cellular phenotype, and changes in cell division timing.

Alternatively, we can look for signatures of developmental processes themselves. Initially, we consider multiple molecular datasets from *Caenorhabditis elegans* (genes, epigenetic markers, and proteins) associated with structural changes and the production of proteins that contribute to cellular remodeling. We then identify a set of genes associated with these proteins, in many cases yielding multiple genes per protein. From these genes, we identify both the gene expression at multiple points of development and the CpG content across the gene sequence. Finally, we identify the homologs of our *C. elegans* protein list in three other taxa: *Drosophila melanogaster*, *Mus Musculus*, and *Saccharomyces cerevisiae*, Other features of genomic evolution between species lead us to a unifying theoretical concept: differentiation waves (Gordon, 1999; Gordon and Gordon, 2016a; Gordon and Gordon, 2019, see also Figures 1 and 2). Differentiation waves (described by differentiation trees) are a reinterpretation of propagating signals that control spatially localized shifts in cell division. Taken together, these concepts allow us to understand the role of collective cell differentiation, tissue formation, and cell lineage in embryogenesis.

**Figure 1.**
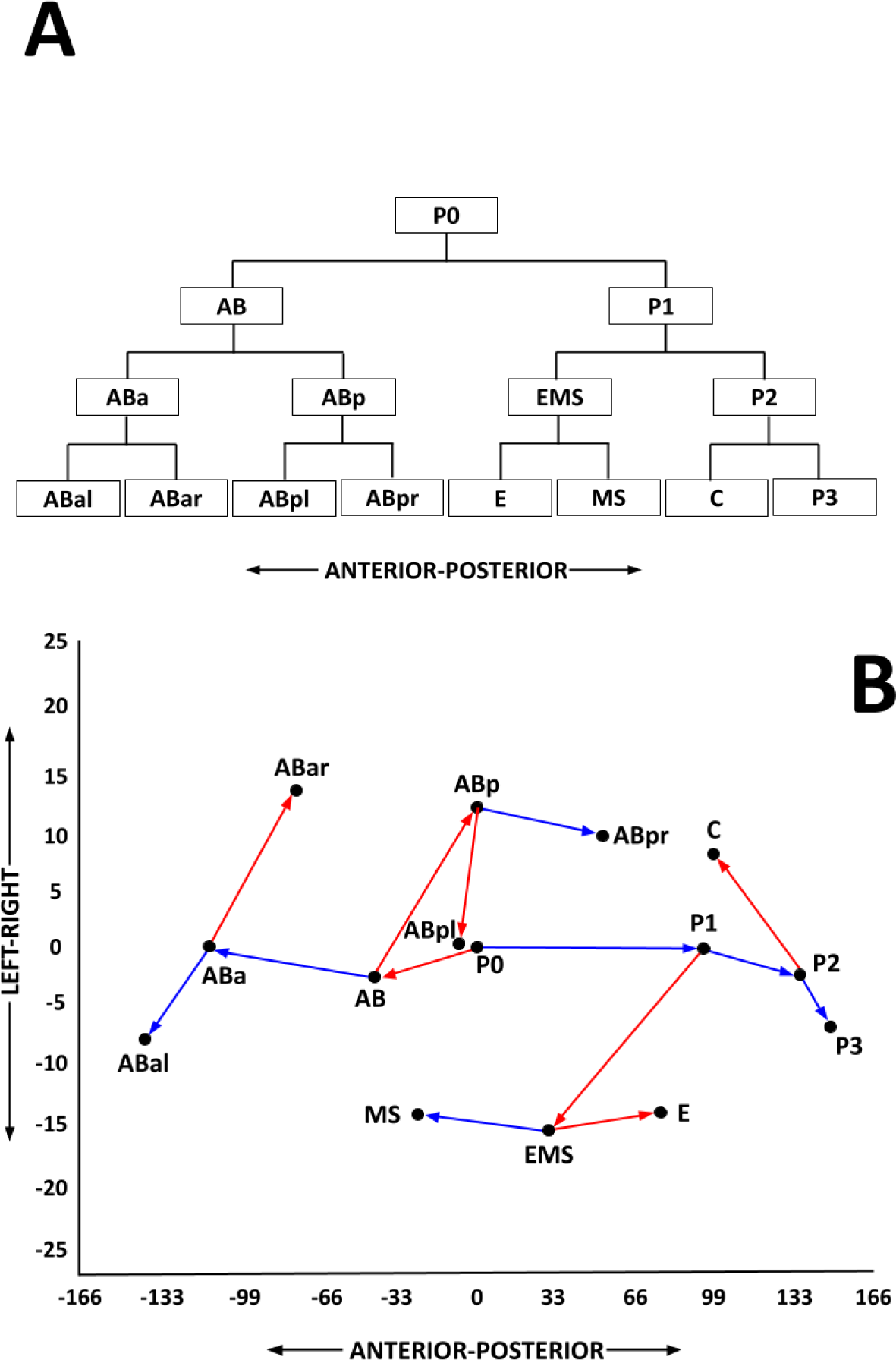
An example of a differentiation tree and corresponding map (Alicea et.al, 2022). A: All founder cells of the *C. elegans* lineage tree are shown in an 8-cell lineage tree with its anatomical orientation (anterior-posterior). B: a 2-D spatial map (anterior-posterior and left-right axes) demonstrating a set of differentiation waves in mosaic development for an 8-cell differentiation tree. Contraction wave-associated branches are shown in red, and expansion wave-associated branches are shown in blue. The position of individual cells, branch points, and their orientation relative to a lineage tree are shown. Coordinates in 1B are normalized cell tracking data from Murray et.al (2012).

**Figure 2.**
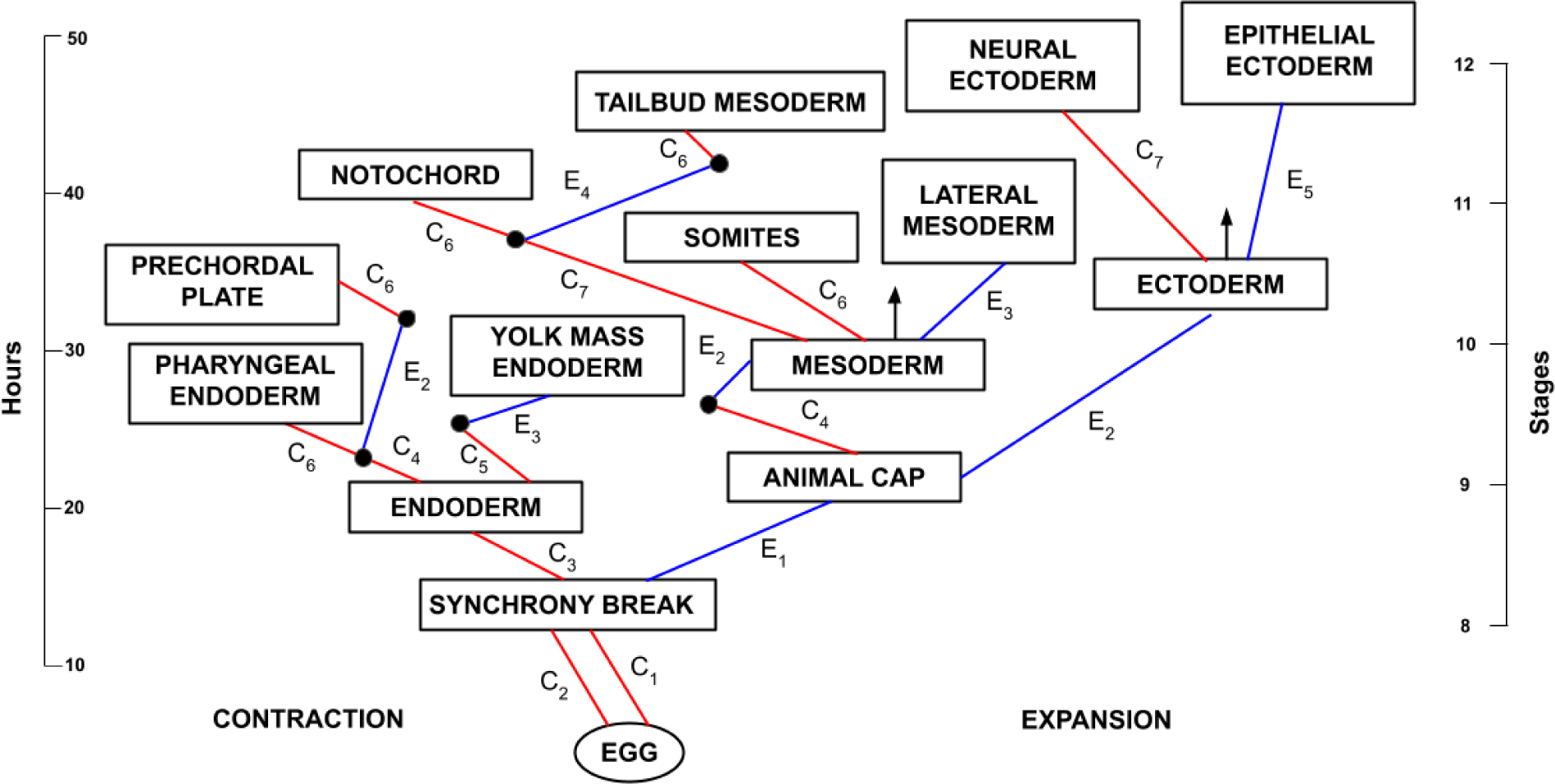
An example of a regulative differentiation tree from the Axolotl (*Ambystoma mexicanum*) from fertilized egg up to notochord formation (developmental stages 8-12 reared at 20°C). All cell types constituting tissue categories are presumptive. Contraction (leftward branch, red) represents a contraction wave, while expansion (rightward branch, blue) represents an expansion wave. *C_n_* and *E_n_* denote the type and number of contraction and expansion waves, respectively. Multiple branches can belong to the same wave (e.g. *E*_2_). Black arrows denote a continuation of an undifferentiated precursor cell population (mesoderm → germline and ectoderm → neural crest cells). Developmental stages and time from fertilization based on Bordzilovskaya et.al (1989). Figure adapted from Gordon and Gordon (2016a), Figure 8.13.

Understanding cellular diversity as the product of multiple differentiation waves requires interactions between phylogenetics, lineage trees, and gene expression in differentiating cells. One of our goals in this paper is to expand our theory of differentiation waves (based largely on cell-level characteristics) with a molecular model of differentiation. Our selected proteins are expressed by gene regulatory systems (see Figure 3), parts of which are included in our data analysis, and produce an output of protein abundance. In our combinatorial model of regulation, a single gene produces transcripts of different loci as shown in Figure 3A. Over the course of evolution, as genes (*g*) are duplicated and functionally diverge, the number of unique interactions between genes in a network increases on the order of *g^n^* (see Methods). This creates a very large space of refined functional outcomes. Yet our bioinformatic analyses and models of interactive contributions (MICs) provide a means to reduce this space in a way that uncovers new aspects of differentiation.

**Figure 3.**
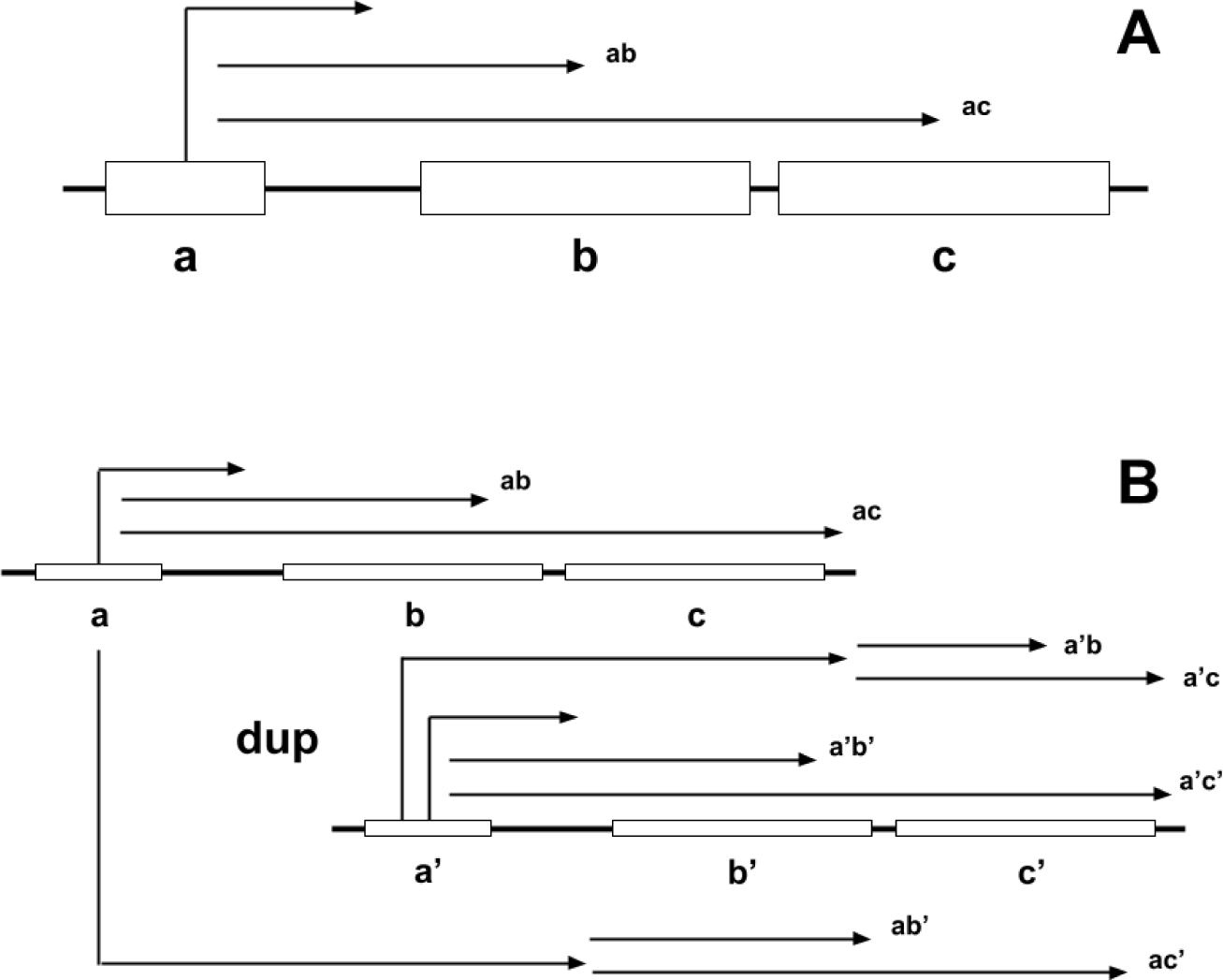
Gene duplication event and the resulting production of gene products. A) the original gene where promoter (a) is expressed to activate either b or c. B) original gene is duplicated, where duplicate promoter (a’) is expressed to activate either b’ or c’. Both the original gene and duplicate gene are involved in co-expression, so that we get regulation that acts both within the same coding region (ab, ac, a’b’, a’c’) and across coding regions (a’b, a’c, ab’, ac’). dup = duplicated genes.

### 1.1 Developmental Differentiation

Across the tree of life, there is a great diversity of developmental types. After eukaryogenesis, an innovation called *continuing differentiation* (Gordon, 1999) that distinguishes most multicellular eukaryotic phenotypes from multicellular prokaryotes (Gordon and Gordon, 2016a). While this allows us to characterize most of eukaryotic development, some systems such as heterocyst differentiation and proliferation in cyanobacteria (Flores and Herrero, 2010), provide evidence of an alternate mechanism: compartmentalized function. As continuing differentiation defines a great many phenotypes, the question arises as to how differentiation trees and their associated waves might map onto the genome. As models of cell and tissue differentiation, differentiation trees are directed acyclic graphs (DAGs), whereas the genomic mechanisms of differentiation waves are hypothesized as being located on multiple chromosomes interacting in a network (Gordon and Gordon, 2016a). Here we will assume that the eukaryotic genome forms a network of candidate genes that produce select proteins that require a mapping of these molecular processes to the differentiation tree.

In organisms such as *C. elegans* or *Ciona intestinalis*, a differentiation tree can be constructed from the deterministic nature of purely mosaic cell differentiation in their embryogenetic phase of development. In Alicea and Gordon (2016), this reconstruction is an axial reorganization of the lineage tree where daughter cells are sorted by cell size (Figure 1). After an initial expansion-contraction event related to the establishment of embryo polarity (AB-P1), each daughter division is sorted left-to-right by their relative size. Larger cells are sorted to the left, and smaller cells to the right. Thus, waves and differentiation are entangled. Each mother-daughter relationship is a single-cell differentiation wave that results in a differentiation event, which allows for continuous differentiation in a sheet of cells.

Local sorting that provides a map to sublineage and ultimately tissue differentiation. In other species such as *Drosophila,* embryogenetic development is partially regulative and so is partially organized using mosaic assumptions, and partially using regulative assumptions. Differentiation trees applied to regulative development, characterized by stochastic differentiation based on cell-cell signaling, are more difficult. In this type of differentiation tree, branches correspond to the origin of different tissue types (Figure 2). Amongst a sheet of cells, cell geometry becomes apically constricted and volumetrically conserved. In Axolotls, a sheet of cells wait for a signal, contract accordingly, and then differentiate (Foe et.al, 1993). Returning to our *Drosophila* example, the imaginal eye furrow exhibits a mix of expansion and contraction events and resembles pull-push dynamics. The differentiation waves themselves are related to mitotic waves that affect the speed of cell proliferation (Foe et.al, 1993), although they are not necessarily correlated.

Within individual cells, there are processes that lock in the differentiated state, and processes that enable the procession of differentiation itself. Based on the transcriptional profiling of developmental cells, there exist molecular events that serve as precursors to major transitions in development. Packer et al. (2019) demonstrates that while a cell’s lineage history is important for transcriptional priming, upon terminal differentiation this priming signature is lost, instead exhibiting the transcriptional signature of the mature cell type. Multilineage priming consists of unique signatures of a sublineage that is then inherited differentially between sister cells and their progeny (Maduro, 2010; Berge et al., 2019; Packer et al., 2019).

Most lineages in *C. elegans* have unique molecular identifiers, which serve as markers of sublineages leading to specific cell types. A drawing of the early *C. elegans* lineage tree up to the 64-cell stage with differentiation-related events is shown in Figure 4. For example, the MS lineage (which leads to mesodermal fates) expresses *ceh-51*, while the MS cell second-order progeny exhibit markers such as *pha-4* (MSaa and MSpa) and *hnd-1* (MSap and MSpp). The levels of expression for these identifiers decrease with developmental time as the progeny become progressively smaller (Packer et al., 2019). By the time the progeny cells terminally differentiate into their adult phenotype, the upregulation of multilineage-specific genes is replaced with the upregulation of genes related to the mature mesodermal phenotype. In *C. elegans*, asymmetric divisions are governed by *pig-1* activity resulting in a variety of timing changes in the lineage tree (Cordes, 2006; Berge et al., 2019). Based on these insights and examples such as *C. elegans* developmental cells exhibiting dual genetic mechanisms for cell identity (blastomere and tissue identity genes, see Maduro, 2010), our differentiation mechanism is likely coupled to broader regulatory networks. On the other hand, proposed mechanisms such as the cell state splitter (Lu et.al, 2012; Gordon and Gordon, 2016b) serve as a molecular mechanism for differentiation linked to cell cycle activity. This along with other mechanisms related to functional changes in the expression of cell cycle genes suggest that there is an alternative way to understand differentiation at the organismal level.

**Figure 4.**
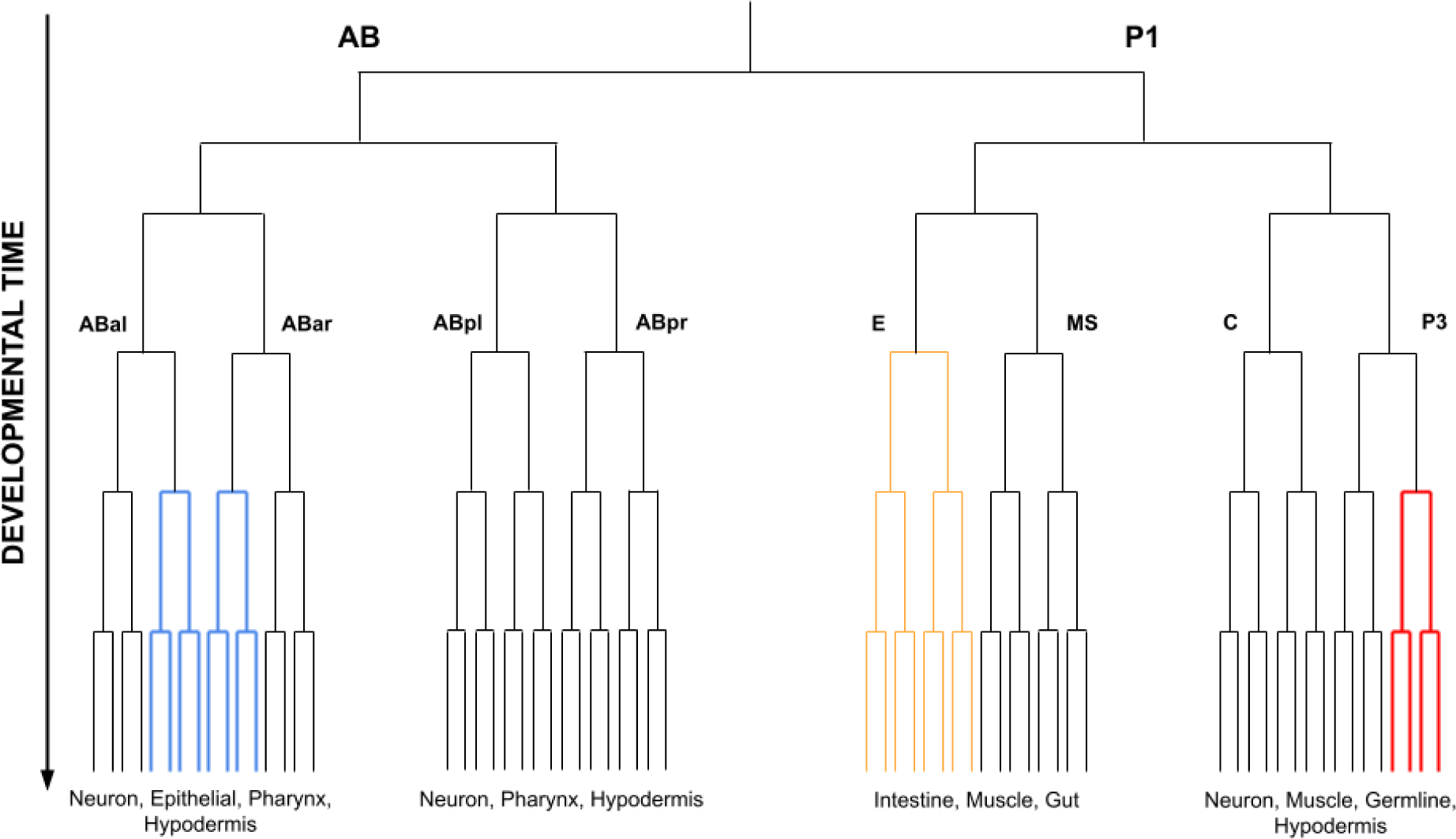
The lineage tree for *C. elegans* up to the 64-cell stage (before variation in division time or asymmetric divisions occur). Blue edges represent parts of the tree where the second Notch has been expressed, red edges represent germline parts of the tree, and green edges represent descendants of the E blastomere that form the gut and intestinal systems (Ewe et al., 2022).

### 1.2 Differentiation Waves as Theoretical Synthesis

Our concept of differentiation waves is tied to a speculative reinterpretation of the developmental biology literature that ties together a number of disparate findings. One candidate mechanism for this is called the differentiation wave (Gordon et.al, 1994; Gordon and Bjorklund, 1996; Nieuwkoop et.al, 1996; Nieuwkoop et.al, 1999). What we refer to as differentiation waves includes diverse embryogenetic phenomena responsible for propagating changes in cell division tempo and the relative synchronization of mitosis. More information on differentiation waves and codes can be found in Alicea and Gordon (2016), Gordon and Gordon (2016b), Gordon and Gordon (2019), Gordon and Stone (2021), and Martin and Gordon (1997). All branches in the differentiation trees shown in Figures 1 and 2 can be interpreted as waves, although they may also be attributable to other causal factors. While differentiation waves precede differentiation itself, differentiation waves generally correspond to the formation of classically-defined tissues formed by changes in the rate of cell division. Specialized structures such as the superficial epithelium of *Xenopus* have also been associated with such oscillations (Nieuwkoop and Koster, 1995). Differentiation waves may not be a universal driver of embryogenesis, but have been observed in a number of model systems and under certain conditions.

Differentiation waves come in two types: contraction and expansion. These relate to the timing of cell division across localized regions of the embryo. Specifically, cell cycle timing is altered in a sublineage of mother and daughter cells, resulting in tissue-level differentiation. Upon moving across the embryo, contraction waves result in shortened intervals of cell division, while expansion waves result in accelerated cell division (Gordon, 1999). Most observations of phenomena that can be classified as differentiation waves have only been observed at the morphological level. Waves, shapes, and movements in the embryo have been tied to the propagation of differentiation waves. As a result, we can only deduce the molecular signatures of differentiation waves from morphogenetic function and protein annotations. Alicea et.al (2018) present a concrete example of how a differentiation wave proceeds in the developing *Drosophila* compound eye. Using high-resolution camera lucida illustrations, we analyzed a differentiating imaginal disc. As the furrow (a marker for coordinated cell division) moves across the imaginal disc at a certain rate, coordinated differentiation occurs with a time delay behind the furrow, yielding the structure of ommatidia that form the compound eye. This also yields a tissue with locally contracted and expanded regions, much like a transformational mesh that characterizes local morphological differences due to growth and diversity of shape (Thompson, 1917; Abzhanov, 2017; Iurato and Igamberdiev, 2020).

The mechanical environment of a developing cell sheet also determines the action of a differentiation wave. When a cell responds to the carrier signal of the differentiation wave, the response of the nucleus (biochemistry) is fundamentally different from that of the cytoskeleton (mechanics). Cell state splitters serve as a pacemaker for differentiation wave-driven differentiation. The process is metastable and can serve to regulate the differentiation process. The splitter consists of an intermediate filament ring that becomes mechanically stuck and checks on the cell’s contraction or expansion state. This sets up the next signal, and so on. Cytoskeletal dynamics carries the differentiation wave, and so like the substrate, current state affects future ones. While we don’t know what triggers and terminates a differentiation wave, candidates include the disassembly of actin, Ca+ flux, and cytoskeleton disassembly. In the blastocoel, a cell tears out of its sheet and becomes migratory for the first time. These cells respond to the differentiation wave signal by forming the neural crest. This involves a cornucopia of biochemical and mechanical signals that lead to pigment cells, neuronal cells, and nerve cells. In general, expansion and contraction occur on a substrate, changes in which can affect the differentiation signal. Contraction provides an example of how differentiation waves act to affect its own mechanical environment. As a contraction wave passes through a population of cells, the geometric connections between cells are changed, which can change the hardness of the substrate. The number of junctions between cells may be of particular importance to tissue stability, an example of which are tri-junctions in epithelial cells (Bosveld et.al, 2018). With a contraction wave, adjoining cells create a hardened space that enhances the substrate in the desired manner. The resulting contraction signal, reinforced by cell-cell and cell-substrate interactions, must then trigger gene expression within individual cells, which then leads to differentiation. While these are separate steps, they are also intimately linked, as discussed in Stone and Gordon (2021).

Synchronous and asynchronous division events are often crucial to understanding the origins of branching in a differentiation tree. Synchronous and asynchronous cell division are often tied to the propagations of different types of differentiation wave, and in turn the degree of morphological differentiation at any one stage of development. Additionally, the rate of cell division provides a means for organismal-level timing mechanisms such as heterochrony (Keyte and Smith, 2014). We can even see how this plays out in organisms without a formal model of development. Green algae undergo a cell division process called multiple fission (Bisova and Zachleder, 2014), where synchronous cell divisions are similar to the cleavage waves observed in Metazoan development. During the life-history of *S. cerevisae*, individual cells make a choice to exhibit different vegetative adhesion strategies, either forming multicellular aggregates or remain in a single-cell phase (Bruckner and Mosch, 2011). Yeast cells also have polarity programs which shape the nature of cell division and result in modifications to the actin cytoskeleton (Chang and Peter, 2003). Whether or not analogous processes outside of formal modes of development can be equivocated with differentiation waves requires comparative analysis.

One way to infer molecular mechanisms responsible for these types of morphological transformation comes from the role cytoskeleton plays in driving the propagation of differentiation waves and the shape of differentiation trees. In general, contraction leads to cleavage arrest, which in turn leads to the inhibition of actin. This prevents a wave of nuclear replication, and requires us to look at targets involving cellular remodeling and generalized processes rather than cell type specific genes. As originally described by Sirakami (1958), cleavage waves allow us to characterize the period and/or rate. Synchronous divisions as consecutive waves. Contraction waves are equivalent to differentiation waves. Ion transport changes in *Xenopus* also provide clues as to proteins of interest.

#### 1.2.1 Eight Examples of Differentiation Waves

Differentiation waves are a reinterpretation of many different types of waves and propagation observed in a variety of model systems from across the tree of life. Many of the events related to differentiation occur during the mid-developmental transition of the phylotypic stage. According to Levin et.al (2016), we should expect genes related to differentiation waves to be regulated differently across phyla. This might transcend the types of development, and indeed our comparisons to *C. elegans* are all from different phyla. Yet we should also expect differences in protein counts to reflect these developmental changes.

Our initial example of the differentiation wave interpretation can be found in the 1-cell embryo. While not universal across development, the fertilization activation wave travels the length of the embryo. As this wave allows for only one sperm to enter the egg, it results in multiple morphological changes. In *Xenopus* embryos (Busa and Nuccitelli, 1985), alteration of the endoplasmic reticulum and the accumulation of actin involves changes in Ca^+2^. In terms of expressed protein, IP3, PKC, and G-proteins. Activation waves have also been observed in Axolotl (Hart et.al, 1992). Other changes occur in early embryogenesis that set up differentiation wave activity. For example, compaction in Mammals pre-blastocoel leads to separation of blastomeres into internal and external cells. The apical constriction process introduces an asymmetry that drives stochastically oriented cell division. This rearrangement is related to cytoskeletal remodeling that minimizes surface area and maximizes cell-cell communication (Lim and Plachta, 2021).

The next example involves a correspondence between the mid-blastula transition and a lengthening of cell cycle timing. This in turn corresponds to an expansion wave, as synchronization of cell division disappears during the mid-blastula transition (Boterenbrood and Naraway, 1986). In terms of direct observations, trains of constriction have been observed in a wide variety of taxa, including fish, newts, crustaceans, and insects. This phenomenon involves a traveling signal transmitted through microfilaments that conform to a contractile ring. Another example of a contraction wave can be found in the ectoderm, where two contraction waves occur in sequence: an initial ectoderm contraction wave followed by a posterior neural plate contraction wave. Coordinated differentiation across the ectoderm is also the basis for the Nieukoop model (Nieukoop and Koster, 1995). According to this model, non-activated ectodermal cells exist ahead of the wave, while the wavefront itself results in activated neuroectoderm along the trailing edge. Taken collectively, these examples contribute to the formation of spatially localized structures.

Surface contraction waves are also a source of differentiation, and thus can be reinterpreted as differentiated waves. One instance of surface contraction waves is observed in the amphibian embryo prior to the mid-blastula transition (Asada-Kubota and Kubota, 1991; Gillis, 1991). The synticial blastoderm of *Drosophila* provides an additional example of waves driving contraction differentiation. In this case, transient actin molecules help along multiple rounds of nuclear division. This results in localized rounds of cell division, during which regions of cells respond in their own manner. Embryogenetic processes such as invagination and evagination also correspond to the rate-limiting of cell division, which in turn may predict the origin of differentiation wave propagation. Invagination results in a slowing down of cell division and decreased mitosis, resulting in the propagation of a contraction wave. Meanwhile, evagination results in a speed up of cell division and increased mitosis, resulting in the propagation of an expansion wave. We can also observe differentiation waves in sea urchin gastrulation, which demonstrates increased mitotic activity. More generally, embryogenetic folding events exhibit localized increases in mitotic activity in a host of species, including flies throughout embryogenesis, in the Mammalian brain, and in the eyeball and primitive streak of the chick.

### 1.3 Theorizing the Molecular Aspects of Differentiation Waves

In terms of scope, the molecular aspects of differentiation waves can be defined as the products of both the genome and proteome across phylogeny and the scope of developmental systems. We have restricted our analysis to examples from three biological contexts: changes in basic polarity and adhesion (*S. cerevisiae*), mosaic development (*D. melanogaster and C. elegans*), and regulative development (*M. musculus*). There is a huge gap between the genome and proteome, particularly for mammalian models. One gene can yield many transcripts and post-translational modifications. As we focus on efforts on *C. elegans*, and then make comparisons at the proteomic level with *D. melanogaster*, *S. cerevisiae*, and *M. Musculus*, we are helped by the well-characterized genome-proteome linkages in *C. elegans* (Schrimpf and Hengartner, 2010). Our strategy is to draw from secondary data on epigenomic signatures, gene expression, and protein abundance associated with a set of genes associated with various aspects of cell differentiation. This type of three-tiered analysis is conducted in *C. elegans*, while further comparisons of protein abundance are made between *C. elegans* and *D. melanogaster*, *S. cerevisiae*, and *M. Musculus*.

We also recognize that there is a gap in understanding between the genome, epigenome, and proteome. Despite having detailed information about whole genome sequencing in model organisms, there are still open questions about how this structure translates to protein expression and phenotypes. At least in the human genome, the number of protein-coding regions is small relative to the large number of functional and non-functional proteins (Salzberg, 2018; Pertea et.al, 2018). The functional significance of post-translational modifications provide an additional layer of complexity (Millan-Zambrano et.al, 2022). Protein diversity in the production of antibodies provides an example of how our protein counts may be inaccurate due to the proliferation of variants. This occurs via a mechanism called switch recombination: the source of this variation is epigenetic, with portions of a small number of coding genes being responsible for the large number of resulting proteins (Alberts et.al, 2002). While this is an extreme case that highlights possible difficulties of mapping genes to proteins, it suggests that we need a deliberate strategy (Suhre et.al, 2021) for taming this complexity.

To overcome some of these limitations, we only focus on the main isoform variants of proteins associated with genes. These associations are derived from and validated by the multitude of available bioinformatics resources: *C. elegans* in particular provides us with good quality community tools. Additionally, our metric for analyzing methylation sites in our target genes provide an expectation for potential combinatorial variation: the greater the number of methylation markers in a gene’s coding region, the greater potential for functional variation (Suelves et.al, 2016). Therefore, we have at least two indicators that can guide us through the combinatorial soup of genome-proteome mapping.

#### 1.3.1 Insights from Single-cell Genomics of *C. elegans*

*C. elegans* embryogenesis has been characterized at the single-cell level using RNAseq. In Tintori et.al (2016), the 16-cell stage (which includes all founder cells) has been characterized. Hashimshony et.al (2015) have also characterized the 2-cell stage, which provides a basis for the first differentiation event in embryogenesis. Packer et.al (2019) have provided information from single cells representing mid-gastrulation embryos, as well as time-points ranging from 210- to 510 minutes of embryonic development. These studies go beyond the traditional genetics and cell-fate specification for particular mechanisms, and provide us with a number of principles. One principle involves the increase in transcriptional diversity over developmental time. In a related fashion, cell fates bifurcate according to a mechanism called multilineage priming. During this process, progenitor cells begin to express genes that are specific to their daughter cell fates. The co-expression of these genes results in all cells nested within a lineage (sharing a common ancestry with the progenitor) to be more transcriptionally similar than distantly related cells. In the context of differentiation waves, we expect there to be fluctuations over time in our candidate genes: decreases over time should lead to contractions in tissue, while increases over time should lead to expansions in tissue. While whole embryo gene expression data is a combination of expansion and contraction waves averaged out over time, individual genes can contribute a signal, which can be taken as an ensemble for proteins of different functions. We should expect a protein like Wnt to have associated genes that contribute a variety of signals to differentiation waves occurring across the embryo. On the other hand, we should expect proteins playing a limited role in developmental processes to possess ensembles that are more homogeneous.

## 2 Proteomics and Epigenomics of Differentiation

### 2.1 Rationale for Proteomic Analysis

We chose a set of candidate proteins that might play a role in driving cell differentiation. These proteins have numerous functions across the embryo, from mediating cell-cell interactions to structural plasticity, and from biochemical signaling to establishing polarity. Of particular interest are kinases and phosphatases, which play an important role in the development of two mosaic developing species: *C. elegans* (Plowman et al., 1999) and *D. melanogaster* (Morrison et al., 2000). In both species, kinases/phosphatases are involved in DNA replication, cell cycle progression, and cell growth/differentiation. But this is not done in a straightforward, linear manner. Kinases and phosphatases amplify and stabilize signaling pathways but turning pathways on and off, respectively, and must exist in a balance for the cell to achieve dynamic regulation.

In addition to our theoretical model, we also conduct an analysis of molecular data from *C. elegans*. First, we define a set of proteins and obtain their abundance across multiple species using secondary data. Then we analyze a set of candidate genes in *C. elegans* for epigenetic signatures of a histone code. The proteomic and epigenomic analyses for *C. elegans* are compared, and those outcomes are generalized to different species representing mosaic, regulative, and a lack of development. We then consider how our candidate genes for their corresponding proteins in *C. elegans* interact, and how those networks provide information about differentiation to cell populations. This is done through considering generic interactions (fully-connected MICs) and ensembles of signals (ordered MICs) that provide information regarding expansion and contraction behaviors.

### 2.2 Epigenomic mechanisms and regulation of cell state

Based on findings from vertebrate cell biology, we suspect that any differentiation switch involves methylation markers that can regulate the activity of gene expression. As epigenetic mechanisms, they are enabled by genomic content. One mechanism is global cell state regulation due to methylation, where histone markers are regulated in a coordinated manner across an individual cell’s nuclear genome. This has been described in stem and reprogrammed cells as bistable molecular systems regulated by self-reinforcing feedback (Gomez-Schiavon and Buchler, 2019). More generally, epigenetic memory can serve to augment the role of transcriptional priming (Cambuli et al., 2014; Madrigal et al., 2023; Osnato et al., 2023). Epigenetic mechanisms provide a memory that is persistent throughout development (Elsherbiny and Dobreva, 2021). We are interested in the bistability demonstrated for differentiating vertebrate cells, and reliant upon CpG dinucleotide signatures (Vincent and Van Seuningen, 2009). In vertebrates, CpG satellites act as promoters in some regions while transcriptional repression results in other parts of the genome. CpG islands, often found in promoter regions, have distinctive patterns of transcriptional initiation and chromatin configuration. In a human stem cell model, it has been found that epigenetic modifications (in the form of CpG methylation) regulate transcriptional activity globally, which in turn primes cell fate (Lunyak and Rosenfeld, 2008; Madrigal et al., 2023).

In mammals, the regulation of methylation is global in nature and occurs in enriched islands that are absent in invertebrates (Allis and Jenuwein, 2016). By contrast, invertebrates have mosaic methylated genomes with alternating methylating and non-methylating domains (Suelvas, 2016), mimicking what is found amongst individual cells in mammalian tissues (Silva et.al, 1993). This should lead to a difference in how differentiation occurs in specific cells of the lineage tree. In *C. elegans* more specifically, Zheng et al. (2013) investigated VC motor neurons and their role in egg laying for two vulval (VC4 and VC5) and non-vulval (VC1-3 and VC6) cells. A mutation of H3K9 methyltransferase MET-2 acts to trimethylate H3K9 and leads to loss of VC cell-specific UNC-4 expression (Schott et al., 2006). Overall, patterns of CpG sites (those repeating over short genomic distances) should enable gene- and protein-specific regulation, thus enabling cell state transitions across the lineage tree (Figure 4). Within the *C. elegans* lineage tree, sublineages express specific genes and proteins well before differentiation (Sulston et al., 1983; Cao et al., 2020). These properties can be characterized using an epigenetic landscape (Moris et al., 2016). Whether asymmetric histone inheritance is a general mechanism during the asymmetric cell division is largely unexplored (Elsherbiny and Dobreva, 2021).

### 2.3 Histone Code and Differentiation

An alternate (or perhaps complementary) hypothesis is that of the histone code: DNA motifs consisting of CG repeats are responsible for triggering differential gene expression that results in cell differentiation. So-called CpG islands are clustered in promoter regions, and control transcription via a bistable mechanism: the gene is poised to switch from on state to an off state, which corresponds to the control of cell fate. *C. elegans* and humans both share highly occupied target (HOT) regions that bind a large number of transcription factors (Turner et al., 2010; Gerstein et al., 2010; Deaton and Bird, 2011; Yip et al., 2012). In *C. elegans* specifically, HOT regions can be occupied by many transcription factors as highly active promoters as well as act in *trans-* (Chen et al., 2019). The *C. elegans* gene CFP-1 acts in a manner like methylation in humans and tends to target genes with a high density of CpG sites (Yoshimura et al., 2019). This suggests that each cell in a lineage tree might have its own specific timing and CpG site dependence, as each terminal differentiation always results in the same cell with a precursor of deep ancestral origin (Memar et al., 2019).

In vertebrates, CpG islands exist as satellite (or long stretches of 2-mer) repeats. In *C. elegans*, CpG repeats are more diffuse, and so require a strategy to capture short regions where multiple CG repeats exist. Candidate genes for which a large and diverse set of repeats exist are genes of high information content, which might correspond to an abundance of its associated protein and thus differentiation potential. We consider this relationship between epigenome and proteome to be the basis for a cell state splitter (Martin and Gordon, 1997; Gordon and Gordon, 2016b). As we also make proteomic comparisons between *C. elegans* and *Drosophila* (as well as *Mus musculus* and *Saccharomyces*), it is worth considering how epigenomic comparative analysis yields similar results due to a high degree of evolutionary conservation in methylation patterns between our two mosaic development species (Ho et.al, 2014).

### 2.4 The Molecular Biology of Differentiation Waves as Computational Models

Using the defined genomic and epigenomic data for *C. elegans*, we can construct two models of interactive contributions (MICs) to better understand the genomic contributions to differentiation wave-related proteins in *C. elegans*. We can look to Genetic Regulatory Networks (GRNs – see Erwin and Davidson, 2009; Walhout, 2011; Wilson et al., 2018; Cussat-Blanc et al., 2019; Grimes et al., 2019; Medweg-Kinney et al., 2020) as inspiration. GRNs provide a block modeling paradigm to infer the functional state of a network where genes regulate each other in their collective expression. As but one example, methylation can act as a repressive force on individual genes, which weights the final output of the model. Given the poor correlative relationship between gene expression and protein abundance, and the difficulty of implementing an interpretable model, we are more interested in the interactions between genes associated with a given protein. Bioinformatics models of genetic interactions and contribution signals to protein expression are informative here (Kelley and Ideker, 2005; Boucher and Jenna, 2013; Cowen et.al, 2017; Gao et.al, 2023), but also do not provide a suitable methodology. As networks, the genes and epigenetic markers that collectively represent a specific protein exhibit a set of interactions amongst themselves (fully-connected MICs) and a collective output that is difficult to interpret simply using measures of expression (ordered MICs).

We thus consider two independent MIC architectures that capture different aspects of regulatory order. Fully-connected MICs capture the potential of gene-gene interactions for a given protein’s associated MIC. The size and expression variety of each MIC serves to define its potential role in developmental order. In some cases, we might expect larger, more variable networks to produce more proteins of importance to the developmental process. By contrast, ordered MICs provide a means to understand the chromosomal ordering and collective contribution of a gene’s output (Medwig-Kinney et.l, 2020) for a given MICs component genes while also extracting component signals related to changes in gene expression over developmental time. The resulting signal mosaics provide cues to potential differentiation waves and the contribution of each protein’s MICs to those waves. In this way, we can build informative MICs unencumbered by neither the specific mechanisms of cell sublineages nor temporal-specific variation such as differentiation in the gut-intestinal system (Ewe et al., 2022).

Genes associated with a certain protein being on different chromosomes tells us something about gene action: genes that are near each other are physically linked with a lower probability of recombination. The concept of synteny suggests that even in the face of duplication events, physically linked genes are more likely to maintain their original function (Simakov et al., 2020). Teichmann and Babu (2004) discuss how duplicated genes allow MICs to grow and gain complexity, and link this to function in *C. elegans*. Functionally-related genes that are also close to one another are more functionally and evolutionarily stable (Szilagyi et al., 2020). In our regulatory computational model, genes being on different chromosomes reduces the probability of upstream promoters being able to regulate downstream genes, as the promoter is less likely to be bound when acting in *trans*. This leads to fewer instances of the a’b, ab’, a’c, and ac’ combinations shown in Figure 3.

#### 2.4.1 From gene expression to proteins

A number of studies point to the joint contribution of transcription factors (TFs) and chromatin modifiers acting as transcriptional cofactors (CFs) to cell differentiation. Both types of regulatory mechanism play a key role in *C. elegans* intestine morphogenesis (Horowitz et.al, 2023). More generally, both TFs and CFs are connected to larger regulatory networks, and involve gene duplications and network rewiring in ways similar to what is described here (Reece-Hoyes et.al 2013). The effects of such regulation can be seen in gene and protein networks. Gene duplications are an important aspect of regulatory networks across species, as Levine and Tijan (2003) estimate that the products of duplicated TFs represent 5-10% of eukaryotic proteomes.

#### 2.4.2 mRNA-protein correlations

Our MICs do not rely on correlations between mRNA and protein abundance, as these have a poor correspondence across genes and context (Maler et.al, 2009). In *Saccharomyces* (Greenbaum et.al, 2003), mRNA aggregate within a cell into pools that exhibit decay, rate-limited protein translation, and transient responses. These result in sublinear (or saturated), linear, and superlinear responses, respectively (Alicea, 2014). To the degree that we can obtain correlations between mRNA and proteins, differentially-expressed gene expression is best (Koussounadis et.al, 2015).

## 3 Bioinformatics Details

Our bioinformatic analyses allow us to explore the molecular underpinnings of differentiation waves in *C. elegans*, and then demonstrate molecular differences at the protein level across different types of development. This is done by reducing the molecular complexity of cells during embryogenesis. To this end, we selected only those candidate genes that have been verifiably associated with a *C. elegans* protein. Calculating the prevalence of CpG sites using Equ [1] for any one gene sequence can tell us about that gene’s contribution to its corresponding protein. Our proteomics data focuses on well-known homologs for cross-species comparison. Taken together, these procedures limit our false positive rate in addition to reducing the chances post-translational modifications obscure the effects of our candidate genes.

### 3.1 Protein Selection

Our candidate proteins are derived from the signaling pathways shown in Figure 8.13 in Gordon and Gordon (2016a), and tend to be evolutionarily conserved (Koh et al., 2016). These include: adenomatous polyposis coli (APC), axin, β-catenin (B-Cat), calcineurin (CaN), Casein Kinase 1 (CK-1), cell division control protein 42 (Cdc42), C-Jun N-terminal kinases (JNK), dishevelled-associated activator of morphogenesis 1 (DAAM1), dishevelled, cytoplasmic phosphoprotein (Dvl), E cadherine (E cad), glycogen synthase kinase 3 (GSK3), inositol 1,4,5-triphosphate (IP3), microtubule associated protein (MAP), mitogen activated protein type K (MAPK), nuclear factor of activated T-cells (NFAT), phosphatidylinositol 4,5-biphosphate (PIP2), phospholipase C (PLC), protein kinase C (PKC), Rac, Rho, Rho associated protein kinase (ROCK), frizzled transmembrane protein (FzR), and *Wnt*.

### 3.2 Blastp and Blastn

A BLAST search was conducted for each protein in our list: using several parameter settings: the maximum target sequence length is 250, the word size is 5, and the *E* threshold: 0.05. The species-specific match was determined using BLAST. For the CpG analysis, genes were selected via literature search (coding genes associated with protein x) and Blast search against the *C. elegans* genome (Sulston and Waterston, 1998).

### 3.3 Comparative Bioinformatics

We assembled a list of proteins for four species: *C. elegans*, *D. melanogaster*, *M. musculus*, and *S. cerevisiae*. These species represent the conditions Mosaic development A (*C. elegans*), Mosaic development B (*D. melanogaster*), Regulative development (*M. musculus*), and basic changes in polarity and adhesion (*S. cerevisiae*). As *S. cerevisiae* exhibits neither regulative or mosaic development, protein abundances should not resemble our other species but should also exhibit elevated abundances of cytoskeleton-related proteins.

### 3.4 Gene expression data for candidate genes in *C. elegans*

We limited our analysis of gene expression to *C. elegans*. Two gene expression datasets were used to estimate gene expression across the progression of a lineage tree. These secondary data describe the state of gene expression in the egg for early (early embryo; up to the 16-cell stage) and middle (late embryo; near hatch and after neurons have differentiated) stages of pre-hatch development (Spencer et.al, 2011), and immediately post-hatch (L1 arrest) for late development (Meeuse et.al, 2020a). This corresponds to the pre-sublineage lineage tree, lineage tree with differentiation tissues, and terminal point of all embryonic cell lineages. Our approach also maps to more general trends in development. Since embryonic development is known to diverge more in early and late embryogenesis than in mid-embryogenesis, our timepoints capture gene expression particular to the *C. elegans* lineage tree. The hourglass model has been validated for gene expression data across phylogenetically diverse *Drosophila* species (Kalinka et.al, 2010), and so should also hold true for our approach to *C. elegans*.

#### 3.4.1 Expression Dataset #1: early and middle development

Gene expression data for *C. elegans* in Table 1 taken from the modENCODE dataset described in Spencer et.al (2011). These data represent whole animal samples with the average expression value for each gene. Our original list of genes are matched with the average gene expression for both early and late embryo conditions (averaged over three replicates for each condition) available via WormMap (https://vanderbilt.edu/wormdoc/wormmap/ <u>Expressed_genes.html</u>). All gene IDs confirmed using WormBase (Harris et.al, 2019). These correspond with our early and middle development timepoints.

**Table 1.**
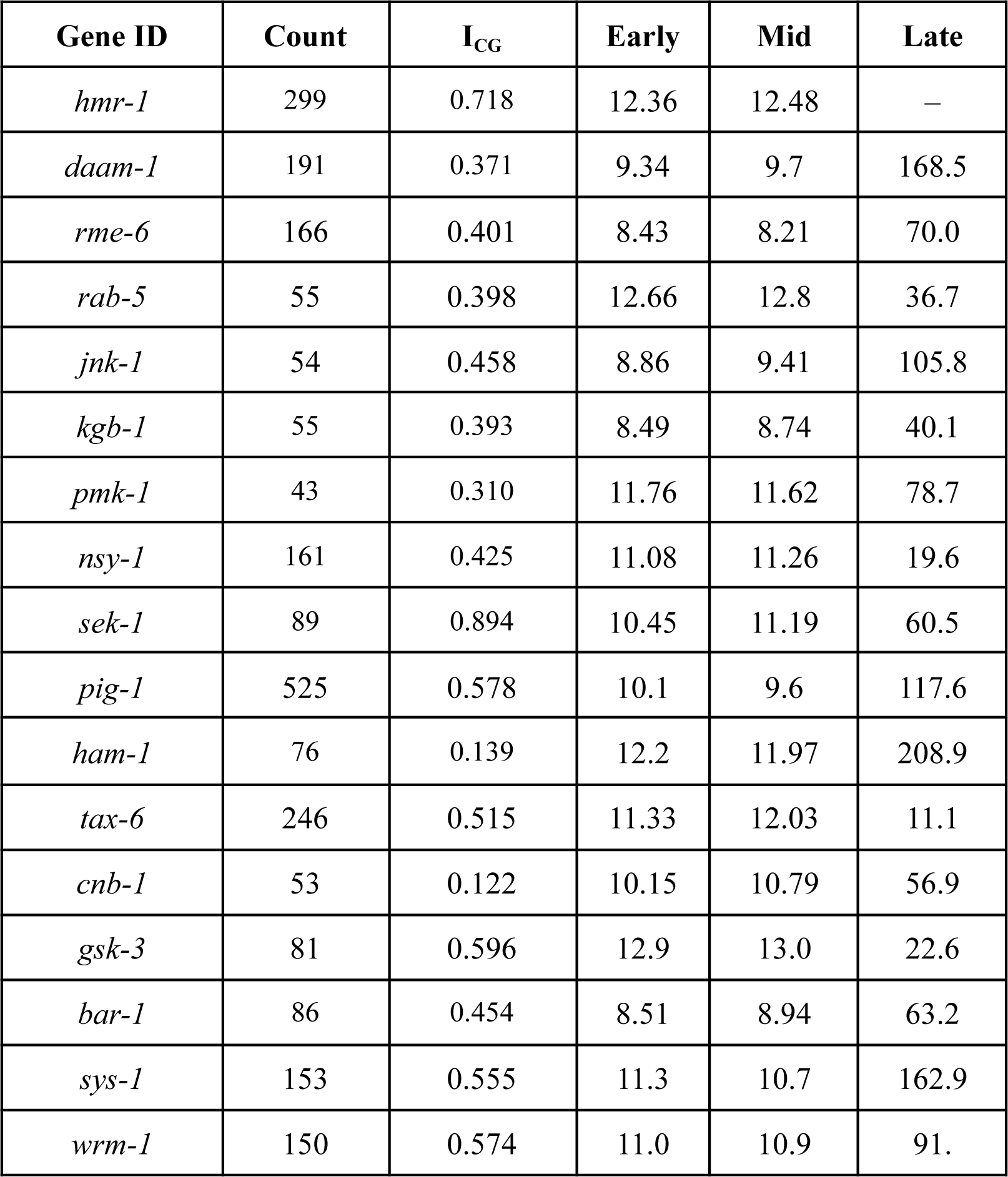
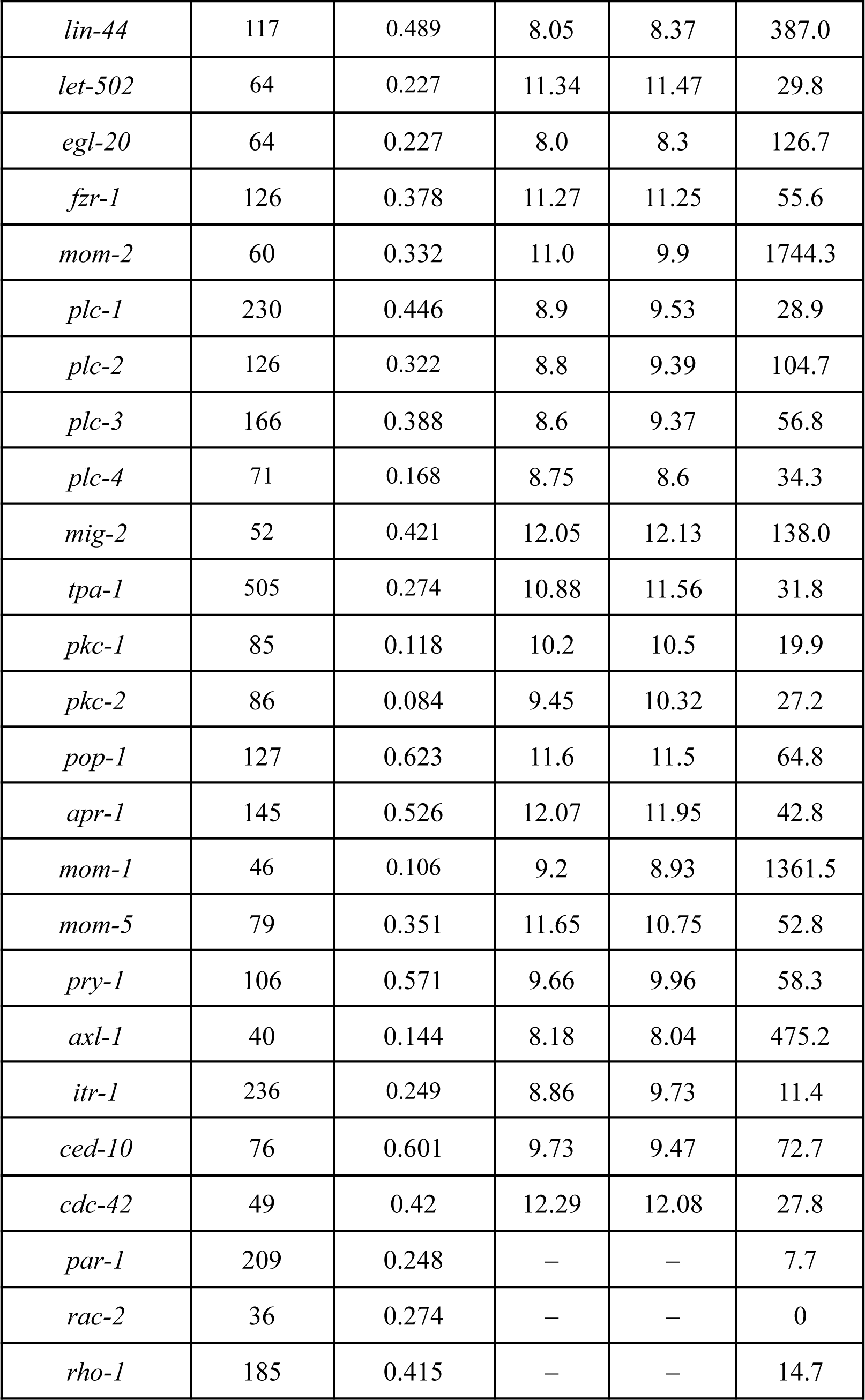
Early embryo (early development) and late embryo (mid-development) average gene expression (whole organism) as compared to CpG data for a subset of gene IDs. Early, middle, and late development are directly comparable except for *hmr-1*, *par-1, rac-2,* and *rho-1* which were not included in both datasets. Normalized RNAseq copy counts from larval worms at hatch (late development) for candidate genes. Normalized using the RPMK approach. Gene list consistent with the WS290 release of WormBase.

#### 3.4.2 Expression Dataset #2: late development

RNAseq data for the *C. elegans* embryo (N2 wildtype) during L1 arrest (larval development) were taken from Meeuse et.al (2020). The gene expression datasets are available on NCBI, Accession number GSE130782. Gene identities were retrieved from Ensembl Metazoa (https://metazoa.ensembl.org/Caenorhabditis_elegans/), consistent with WormBase WS282. Data was collected using an Illumina HiSeq 2500 using *C. elegans* oligonucleotides.

A list of RNAseq average normalized gene expression values is shown as the late development column of Table 1. RNAseq values are normalized by taking the average of three genes: *rps-2*, *rps-4*, and *rps-23*. Normalization is consistent with the RPKM approach to normalizing expression values (Tao et.al, 2020). These correspond with our late development timepoints.

### 3.5 Gene and Protein Sequences

#### 3.5.1 *C. elegans* Candidate Genes

Candidate genes identities for *C. elegans* proteins are taken from the following references: *Wnt* and β-cat (Jackson and Eisenmann, 2012), CK-1 (Guillen et al., 2020), E-cad (Klompstra et al., 2015), Ras (Sato et al., 2005), PKC (Li et al., 2018), CaN (Dwivedi et al., 2009), Rho (Hahmann and Schroeter, 2010), and Rac (Lundquist et al., 2001).

#### 3.5.2 Chromosomal Distribution of Candidate Genes

Figure 5 shows the distribution of candidate genes across all chromosomes in the *C. elegans* genome. While the genes are widely distributed, autosomes I and IV provide greater than 10 genes each. Sex chromosome X provides 7 genes.

**Figure 5.**
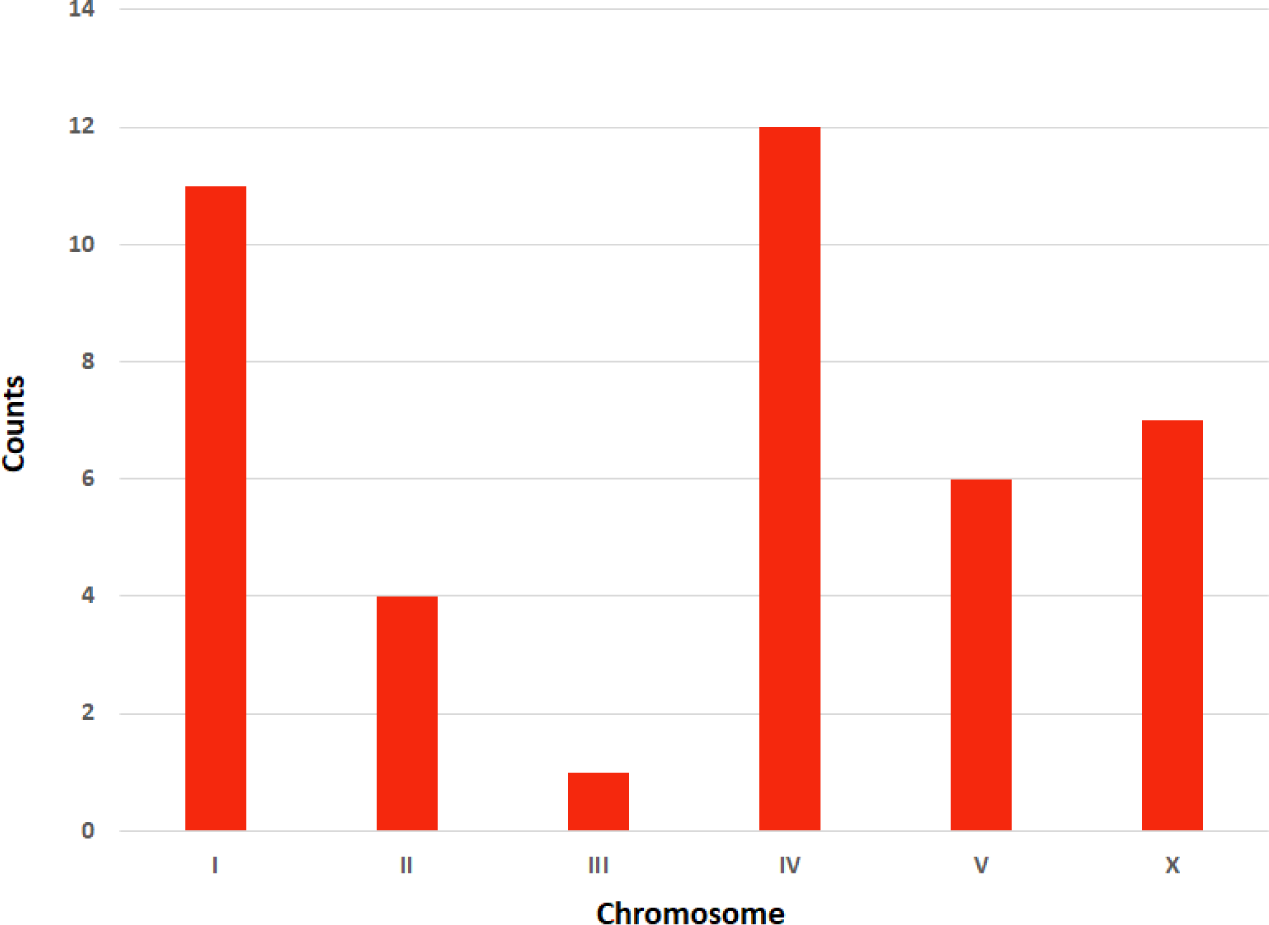
Distribution of candidate coding genes across the *C. elegans* genome by chromosome.

#### 3.5.3 *C. elegans* gene identities

The following identities were used, and NCBI accession numbers represent the source of the DNA sequence: *bar-1* (NCBI: NM_076805.8), *cdc-42* (NCBI: NM_063197), *cnb-1* (NCBI: NC_003283.11), *cwn-2* (NCBI: NM_069421.6), *kgb-1* (NCBI: NM_067521.8), *jnk-1* (NM_001392324.1), *hmr-1* (NCBI: NM_00139294.1), *ham-1* (NCBI: NM_070000), *daam-1* (NCBI: NM_070231.6), *egl-20* (NC_003282), *fzr-1* (NCBI: 003280), *gsk-3* (NCBI: NM_001392959.1), *let-502* ((NCBI: NM_070000), *lin-44* ((NCBI: NC_003279), *mom-1* (NM_076653.6), *mom-2* (NCBI: NM_072753), *mom-5* (NM_060234.7), *nsy-1* ((NCBI: NM_001383826.1), *par-1* (NCBI: NC_003283.11), *pig-1* (NCBI: CP_11639.1, contig), *pkc-1* (NCBI: NM_001269466, contig), *pkc-2* (NCBI: NM_001129650), *pkc-3* (NCBI: NC_003280), *pmk-1* (NCBI: NC_003280), *rab-5* (NCBI: NM_060080.7), *rho-1* (NCBI: NC_003282.8), *rme-6* (NCBI: NM_001392724.1), *sek-1* (NCBI: NM_076921.8), *sys-1* (NCBI: NC_003279.8), *tax-6* (NCBI: NC_003282), *tpa-1* (NCBI: NC_003283), and *wrm-1* (NCBI: NC_003281), *let-502* (NM_059039), *plc-1* (NM_001136455.4), *plc-2* (NM_074351.6), *plc-3* (NM_063804.5), *plc-4* (NM_068812.5), *tpa-1* (NC_003283), *pop-1* (NM_0586526), *apr-1* (NM_001026376.6), *pry-1* (NM_061073), *axl-1* (NM_001047206), *itr-1* (NM_001028001.5), *ced-10* (NM_067962.7), *mig-2* (U82288.1), and *rac-2* (NM_001047496.1).

#### 3.5.4 *C. elegans* protein identities

The following identities were used, and NCBI accession numbers represent the source of the DNA sequence: APC (Cell Adhesion; NCBI: SFQ94271.1), AXIN (*Wnt* signalling; NCBI: ABA28304.1), β-Catenin (Cell-cell adhesion; NCBI: CCE71280.1), CaN (calcineurin, immune system; NCBI: O02039.1), CK-1 (casein kinase, *Wnt* signalling; NCBI: AAP06180.1), Cdc42 (cytoskeletal rearrangements; NCBI: NP_495598.1), JNK (stress pathway; NCBI: BAA82640.1), Ecad (cell-cell adhesion; NCBI: CCD72204.1), GSK3 (embryonic differentiation; NCBI: CAA22311.1), IP3 (intracellular Ca concentrations, growth factor; NCBI: CAB45863.1), MAPK (related to JNK; NCBI: CCD61895.1), NFAT (development of muscles, nervous system), PLC (Phosphorylation; NCBI: VTW47462.1), PKC (Phosphrylation; NCBI: VTW47465.1), Rac (modulation of cytoskeleton, various cellular functions; NCBI: CAB04593.1), Rho (neuronal survival, axon regeneration; NCBI: CCD69972.1), ROCK (cytokinesis; NCBI: P92199.1), FzR (cell cycle progression; NCBI: ABA18181.1), *Wnt* (morphogen; NCBI: AAA64847.1), and MELK (maternal leucine zipper kinase protein).

#### 3.5.5 *D. melanogaster* protein identities

The following identities were used, and NCBI accession numbers represent the source of the protein sequence: APC2 (NCBI: AAB41404.1), Axin (NCBI: AAF21293.1), CaN (NCBI: NP_524600.3), CK-1 (NCBI: CAA64358.1), Cdc42-related (NCBI: NP_001245762.1), JNK (NCBI: AAB48381.1), DAAM-1 (NCBI: NP_726723.2), Dvl (NCBI: NP_511118.2), IP3 (NCBI: NP_001287180.1), MAP (NCBI: CAB55772.1), MAPK (NCBI: NP_001287635), NFAT (NCBI: NP_511147.2), PLC (NCBI: AAA28724.1), PKC (NCBI: AAA28817.1), Rho (NCBI: CAA36692.1), ROCK (NCBI: NP_001285337), and FzR (NCBI: NP_001261836.1).

#### 3.5.6 *S. cerevisiae* protein identities

The following identities were used, and NCBI accession numbers represent the source of the protein sequence: APC (NCBI: KAF1901282.1), CaN (NCBI: CAA86718.1), CK-1 (NCBI: CAA42896.1), Cdc42 (NCBI: GFP65687.1), GSK3 (NCBI: KAF1903835.1), MAP (NCBI: GAX68832.1), MAPK (NCBI: GHM91332.1), PIP2 (NCBI: CAA99692.1), PLC (NCBI: AAA99927.1), and PKC (NCBI: GAX69463.1).

#### 3.5.7 *M. musculus* protein identities

The following identities were used, and NCBI accession numbers represent the source of the protein sequence: APC (NCBI: XP_030106138.1), Axin (NCBI: AAC53285.1), β-catenin (NCBI: AAA37280.1), CaN (NCBI: AAA37432.1), Cdc42 (NCBI: NP_001407060.1), MAPK (NCBI: CAC88132.1), DAAM1 (NCBI: XP_036013190.1), GSK3 (NCBI: NP_001026837), IP3 (NCBI: EDK99409.1), Fam82a1 (NCBI: FAA00419.1), MAPK (NCBI: AAH58719.1), NFAT5 (NCBI: NP_061293.2), PLC (NCBI: AAH65091.1), PKC (NCBI: BAA14288.1), Rhodopsin (NCBI: XP_006505923.1), ROCK ½ (NCBI: XP_036013163.1), and Frizzled (NCBI: EDL36031.1).

### 3.6 CpG Analysis in *C. elegans*

To find the number and frequency of CpG sites associated with our candidate proteins, we determined coding genes associated in the *C. elegans* genome with each protein. Table 2 provides a mapping between candidate coding genes and proteins, in addition to the chromosome on which they reside.

**Table 2.** *C. elegans* Gene ID with associated protein and Chromosome. Source: WormBase.

| Gene ID | Protein | Chromosome |
| --- | --- | --- |
| *hmr-1* | Ecad | I |
| *daam-1* | Daam | V |
| *rme-6* | CK-1 | X |
| *rab-5* | CK-1 | I |
| *jnk-1* | JNK | IV |
| *kgb-1* | JNK | IV |
| *pmk-1* | MAPK | IV |
| *nsy-1* | MAPK | II |
| *sek-1* | MAPK | X |
| *pig-1* | MELK | IV |
| *par-1* | MELK | V |
| *ham-1* | MELK | IV |
| *tax-6* | CaN | IV |
| *cnb-1* | CaN | V |
| *fzr-1* | FzR | II |
| *gsk-3* | GSK-3 | I |
| *bar-1* | β-cat | X |
| *sys-1* | β-cat | I |
| *wrm-1* | β-cat | III |
| *lin-44* | Wnt | I |
| *egl-20* | Wnt | IV |
| *mom-2* | Wnt | V |
| *cdc-42* | CDC42 | II |
| *let-502* | ROCK | I |
| *plc-1* | PLC | X |
| *plc-2* | PLC | V |
| *plc-3* | PLC | II |
| *plc-4* | PLC | IV |
| *tpa-1* | PKC | IV |
| *pkc-1* | PKC | V |
| *pkc-2* | PKC | X |
| *pop-1* | APC | I |
| *apr-1* | APC | I |
| *mom-1* | APC | X |
| *mom-5* | APC | I |
| *pry-1* | AXIN | I |
| *axl-1* | AXIN | I |
| *itr-1* | IP3 | IV |
| *ced-10* | Rac | IV |
| *mig-2* | Rac | X |
| *rac-2* | Rac | IV |

### 3.7 Definitions

#### 3.7.1 Definition of a gene

A gene is defined as a coding region associated with an interval matching the gene descriptions (an interval between two genomic locations) defined by Sulston and Waterston (1998). Genes sequences are either derived from mRNA associated with that interval, or an intron representing part or all the intervals. All intronic regions are found by conducting a Blastn against the *C. elegans* genome, and are confirmed using the *C. elegans* Genome Data Viewer (https://www.ncbi.nlm.nih.gov/genome/gdv/browser/genome/?id=GCF_000002985.6).

#### 3.7.2 Protein count definitions

Protein counts were conducted using the Blastp function through NCBI’s web interface. From our list of protein identities, a sequence with the highest number of amino acids is retrieved from the NCBI database. A Blast search is conducted for each protein sequence for a 250 maximum target sequence length and the BLOSUM62 alignment algorithm. All unique results with the same identity are then counted, which excludes isoform diversity. A count was considered to be zero when no BLAST match was found within the taxonomic category of interest.

#### 3.7.3 Information Measure

For any given 12 nucleotide-wide window, we calculate all combinations of CpG repeat within that window. While this window can include a contiguous repeat, it can also include multiple, non-contiguous repeats as well. The diversity of repeat types across each gene sequence are calculated as our measure *I_CG_* The *I_CG_* for a single gene is defined mathematically as

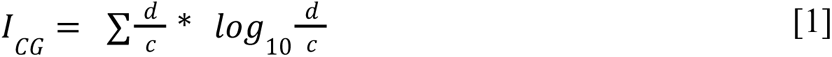

Where *d* is the density of a 12-mer window, defined as the number of repeats and the distance between those repeats (e.g. 3 CG repeats in an 11 base interval), and *c* is the count of all occurrences of CG sites across the gene sequence. Correspondence between *I_CG_* and average gene expression in *C. elegans* is shown in Figure 6. Average gene expression outliers have a low *I*_*CG*_ value, while there is no correlation across candidate genes.

**Figure 6.**
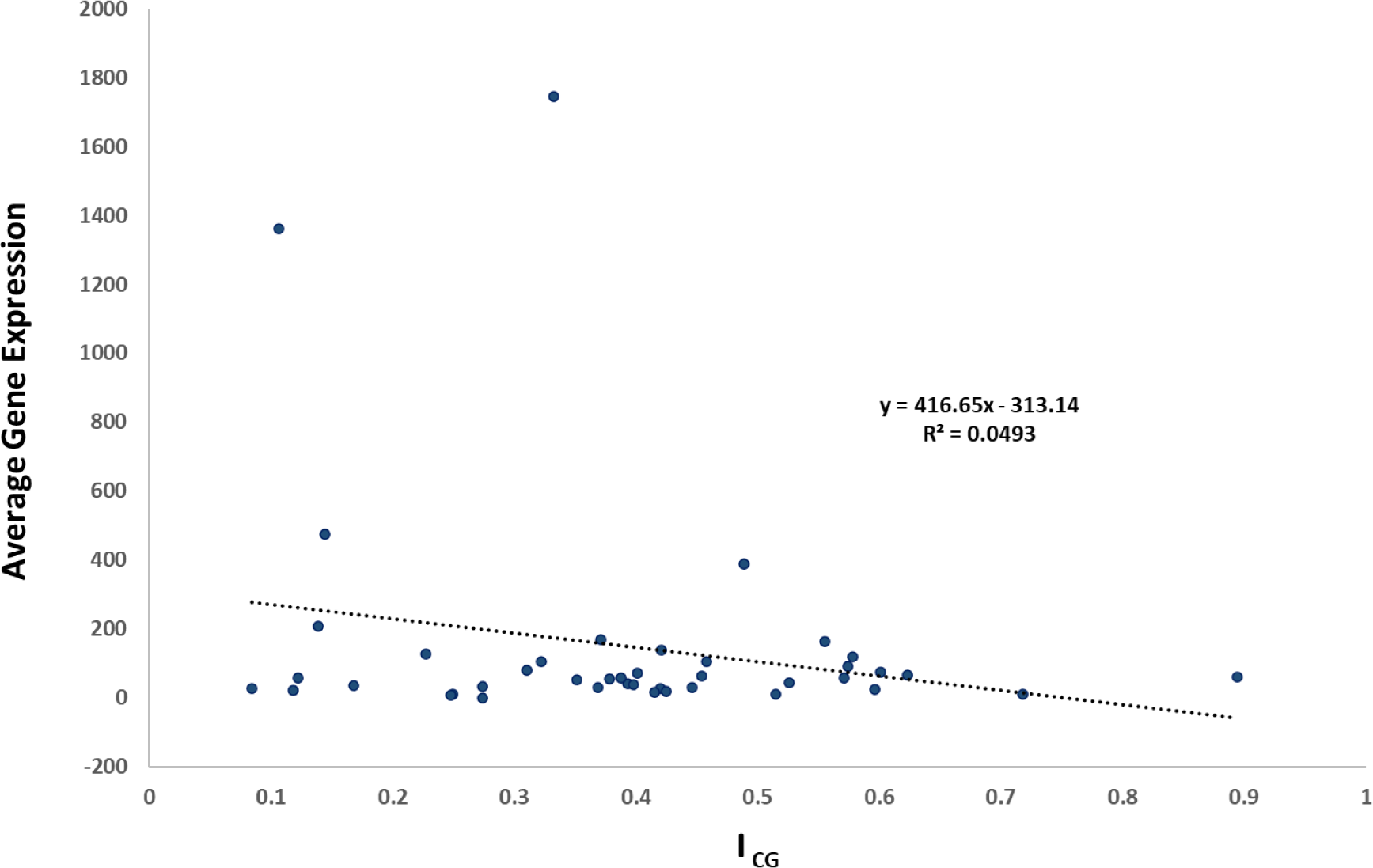
A comparison of *I_CG_* versus average gene expression (normalized) for all genes in the late development gene expression data (*C. elegans*) shown in Table 1.

#### 3.7.4 Division Type

The specific example of mosaic development featured in this paper (*C. elegans*) exhibits three types of cell division: *symmetric divisions*, *asymmetric divisions*, and *terminal differentiations*. While *transitory differentiation* (differentiation to intermediate cell types) occurs post-embryonically, they are not significant in pre-hatch development.

### 3.8 Regulatory Computational Model

Our regulatory computational model incorporates gene expression, epigenomic information (*I_CG_*) for the corresponding gene, and protein abundance for all associated genes to reveal what is required from the intermediate components to produce a linear output. We are not identifying nor predicting interactions between genes. Rather, we want to fit together the relationship between our target genes as to how they produce a potential state change. Multiple genes can correspond to single proteins, thus influencing the pathways through the network. Although we are not making a direct connection to protein abundance with our MIC models, the effects of genomic regulation are non-deterministic with respect to the proteome. A stochastic process of combinatorial generation of alternative transcripts, which can result in a potentially large proteomic space. This is indeed the state of affairs in cancer and immune cells (Bernard et.al, 2022). While our MIC represents coded components, we could also incorporate stochastic components (which we will revisit in Section 4.5).

#### 3.8.1 Fully-connected MIC

A fully connected network is defined by the determinant of the connectivity matrix A. We take the absolute value of the determinant to yield ||A||. The log transform of ||A|| is used to make meaningful comparisons. ||A|| is calculated using the detr( ) function in SciLab 2024.0.0. A fully-connected network is of size *g^2^*, where *g* is the number of independent genes associated with a single protein. We produce 17 networks, each containing the interactions between genes associated with each protein, and summarized by the determinant ||A|| (see Appendix A). In cases where only a single gene is associated with the protein, the singular RNA copy number is identical to determinant ||A||. The ICG measure for each gene is used to weight each interaction, which in turn generates matrix *W_ij_*. This provides a transformed matrix determinant (||A||’) determined by the following equation

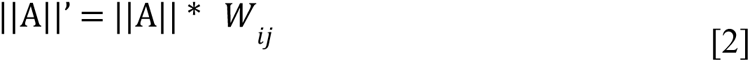

#### 3.8.2 Ordered MIC

An ordered MIC utilizes the early development, mid-development, and ICG measure derived from sequence data for each gene in our 17 networks, and orders the connectivity by position of each element in the genome. Starting at the beginning of Chromosome I (1bp location) and proceeding through Chromosome X, the ordering of genes proceeds by chromosome and location as determined by NCBI’s Genome Data Viewer for *C. elegans* (https://www.ncbi.nih.gov/gemone/gdv/browser/genome). We produce 17 networks for each gene, and the output of each network is defined by four compartments (see Appendix B). An ordered MIC for a given protein is defined as

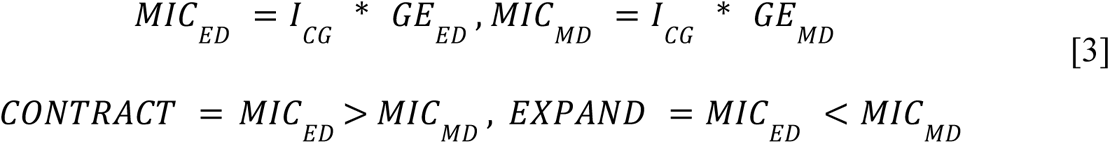

where *MIC_ED_* is the early development ordered MIC for a given gene, *MIC_ED_* is the late development ordered MIC for a given gene, *I_CG_* is the information for CpG sites in a given gene, *GE_ED_* is the early development gene expression value for a given gene, and *GE_ED_* is the mid-development gene expression value for a given gene. The contraction and expansion criteria are also presented as decreases and increases in gene expression over time, respectively. The *I_CG_*value is used to weight the average gene expression so that *I_CG_*values closer to 1.0 are closer to the full expression potential of a gene.

### 3.9 Mechanisms for Transitions to Differentiation

The transition to differentiation involves genes and their duplicates (regulatory computational model) that lead to the expression of proteins. In some cases, this relationship is straightforward. For example, *plc-1*, *plc-2*, *plc-3*, and *plc-4* all contribute to the PLC protein. In other cases, the genes associated with a protein are not physical duplicates but seem to share a function. *rab-5* and *rme-6* are functionally associated with CK-1, and *pry-1* and *axl-1* share the same association with Axin. In both cases, duplicates from two different families have converged upon association with a single protein. Some of these associated genes are on the same chromosome (pry-1 and axl-1), while others are on different chromosomes (*plc-1*, *plc-2*, *plc-3*, and *plc-4*).

*C. elegans* provides the ideal model for converting the lineage tree to a differentiation tree to a differentiation code at the cellular level. The *C. elegans* embryo produces 671 cells before hatching, all of which have a deterministic fate (Sulston et al., 1983). As 113 of these cells undergo apoptosis, this leads to 556 somatic and 2 germ cell precursors upon hatching (Kipreos, 2005). These opportunities for asymmetric division produce several gaps in the differentiation code that propagate to higher levels of the differentiation tree. But what is the specific molecular mechanism that contributes to asymmetric divisions and terminal divisions? Given the differentiation tree of an organism, we anticipate that the number of DNA motifs in a genome should result in binary constraints with respect to the differentiation tree, characterized by the number of cell types that occur during embryogenesis (discussed in Bonner, 1988; Gordon, 1999).

Repeating motifs in DNA are well known and studied (D’Haeseleer, 2006), and usually far exceed the number of cell types (Treangen and Salzberg, 2011). Motifs generally carry information or provide the basis for a transcriptional mechanism. We use a specific measure of CpG islands within a coding region of a gene (information) that summarizes the pattern of repeats. The first step of differentiation likely involves the opening of chromatin in some regions and closing in others (Chen and Dent, 2014), along with post-transcriptional control of gene expression (Corley et.al, 2020; Turner and Diaz-Munoz, 2018). As a consequence, the greater the CpG information in the coding region, the more protein activity (embodied in a larger count) we expect to find.

## 4 Results

This section will begin by showing the genomic-epigenomic-proteomic linkages for our selected differentiation mechanism candidates in *C. elegans*. This will include an epigenomic analysis and a proteomic analysis. The epigenomic analysis examines the protein counts and their epigenomic enrichment, in addition to what is revealed by our previously defined epigenomic information (*I* ) measure. Proteomics contributions to our analysis suggest that the results from *C. elegans* generalize to our other mosaic development taxon (*Drosophila*), but not as much to our other biological comparisons. We then map these results to our differentiation wave hypothesis. This includes a mechanism for differentiation in mosaic development, MIC modeling using both a fully-connected MIC and an ordered MIC. The final result is an exploration of how the molecular indicators of differentiation map to differentiation trees, and how differentiation trees can change over the course of evolution.

### 4.1 Epigenomic Analysis in *C. elegans*

#### 4.1.1 Protein counts and enrichment of the epigenome

Our analysis of CpG sites for our 42 candidate coding genes is shown in Table 1 and Figure 7. Figure 7 provides a comparison of *C. elegans* epigenetic and proteomic data, which may provide correspondence between the histone code hypothesis and picking candidate proteins from their annotations. Various comparisons of averaged information (*I*_*CG*_) and direct CpG counts yield few strong trends in the data. In Figure 7B, except for MELK and PKC, there is a negative trend between CpG sites and proteins. In general, as the prevalence of a certain protein increases, the number of CpG sites amongst the candidate genes decreases. MELK and PKC show the opposite pattern, with high protein and CpG site counts. Figure 7D shows the relationship of Figure 7B for all candidate genes instead of for averages, and the result shows that for the most part, a low protein count corresponds to a low CpG site count. Most points cluster in the lower left-hand portion of the graph. Figure 7C compares our information measure with an averaged count of CpG sites. Only four proteins score above the 50^th^ percentile for both information and CpG sites count: APC, β-cat, Ecad, and MAPK. This tells us that the candidate genes for each of these proteins both possess many CpG sites and have many varied repeat motifs.

**Figure 7.**
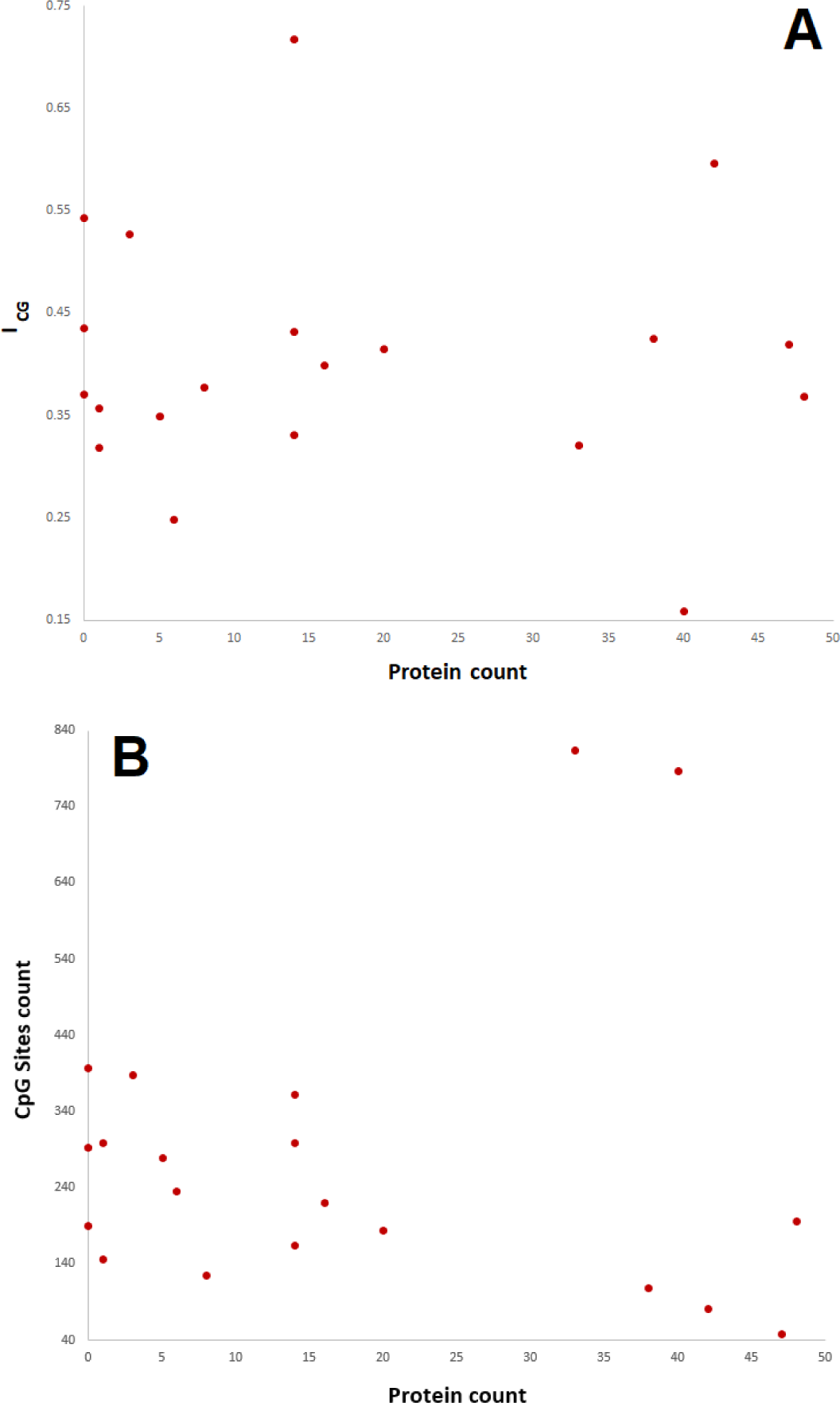
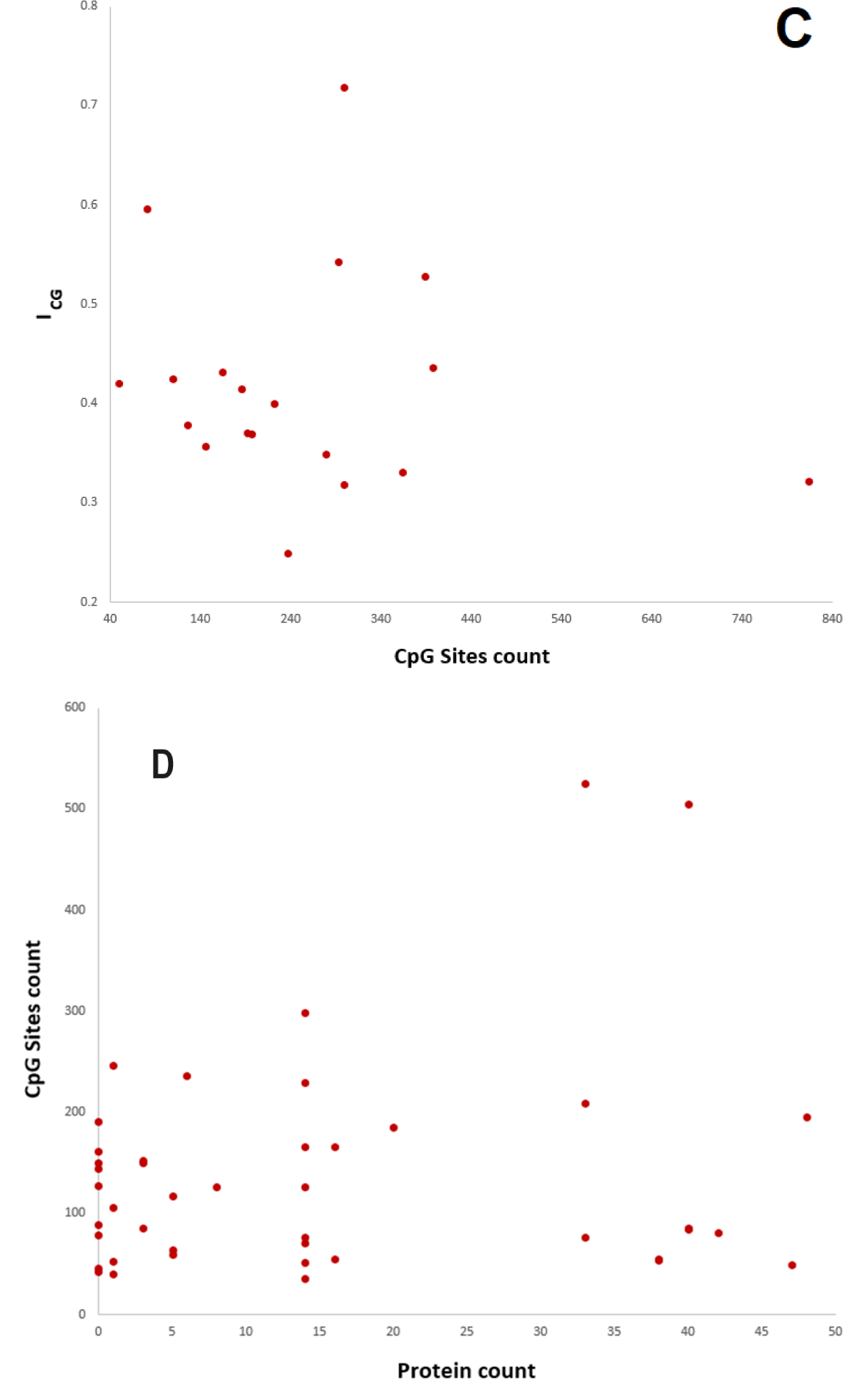
Bivariate relationship between epigenomic and proteomic measures. A) Protein counts versus *I_CG_* B) Protein count versus CpG Sites count. C) CpG Sites count versus *I_CG_*. NFAT is excluded as *C. elegans* has lost NFAT during evolution (Irazoqui et al., 2010). D) A restatement of Figure 3B for every candidate gene.

#### 4.1.2 Epigenomic information and the enrichment of gene expression

The gene expression data in Tables 1 and 2 can be normalized by the *I*_*CG*_ measure for both early and middle development. Figure 8 is a comparison between early and mid-developmental dataset (change in gene expression for genes across embryogenesis).

**Figure 8.**
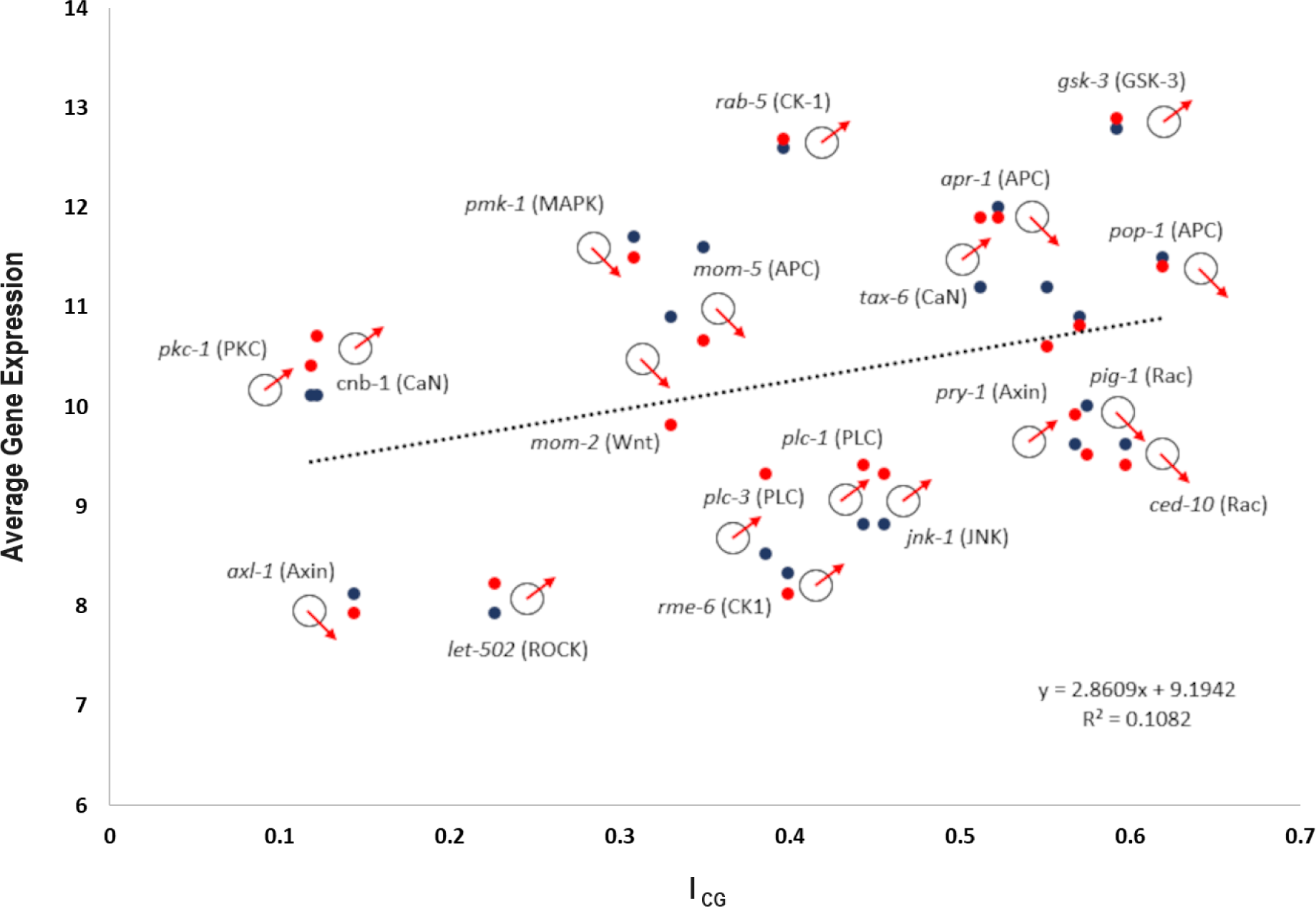
A comparison of *I_CG_* versus average gene expression (microarray) for all genes in Tables 1 and 2. Blue: early development dataset. Red: mid-development dataset (late embryogenesis). Pairs of gene expression values are vertically stacked (e.g. blue above red indicates a loss of gene expression across embryogenesis). Each pair is labeled with a gene name and corresponding protein in parentheses. Circle with red arrows oriented upward and downward represent positive and negative signals from each gene in the ordered MIC (see Figure 12 and Appendix B). The direction determined by the ordered MIC may not match the stacking order of data points for a single gene.

### 4.2 Proteomic Analysis in *C. elegans*

Our epigenomic analysis reveals six proteins of interest in *C. elegans*: APC, β-cat, Ecad, MAPK, MELK, and PKC. These proteins might be important drivers of differentiation, and thus trigger or be changed as a result of the cell state splitter. The first comparison made is intra-specific (*C. elegans*). Table 3 shows the counts for all protein identities while also allowing us to compare protein counts with the epigenomic analysis. The first four (APC, β-cat, Ecad, and MAPK) demonstrate the negative relationship between epigenomic and proteomic mechanisms, as only β-cat (3) and Ecad (14) are expressed to a significant degree in *C. elegans* (Jackson and Eisenmann, 2015; Klompstra et al., 2015). Our other two proteins of interest (MELK and PKC) are robustly represented (counts of 33 and 40, respectively) in *C. elegans*.

**Table 3.** CpG sites for all candidate coding genes with corresponding protein. Information (**I_CG_**) measured by *H*-value (see Methods). *wrm-1* gene codes for proteins β-cat and APC. PLC genes are also associated with IP3 protein function (Baylis and Vazquez-Manrique, 2012).

| Gene ID | Protein | Count | $I_{CG}$ |
| --- | --- | --- | --- |
| *hmr-1* | Ecad | 299 | 0.718 |
| *daam-1* | Daam | 191 | 0.371 |
| *rme-6* | CK-1 | 166 | 0.401 |
| *rab-5* | CK-1 | 55 | 0.398 |
| *jnk-1* | JNK | 54 | 0.458 |
| *kgb-1* | JNK | 55 | 0.393 |
| *pmk-1* | MAPK | 43 | 0.310 |
| *nsy-1* | MAPK | 161 | 0.425 |
| *sek-1* | MAPK | 89 | 0.894 |
| *pig-1* | MELK | 525 | 0.578 |
| *par-1* | MELK | 209 | 0.248 |
| *ham-1* | MELK | 76 | 0.139 |
| *tax-6* | CaN | 246 | 0.515 |
| *cnb-1* | CaN | 53 | 0.122 |
| *rho-1* | Rho | 185 | 0.415 |
| *fzr-1* | FzR | 126 | 0.378 |
| *gsk-3* | GSK-3 | 81 | 0.596 |
| *bar-1* | β-cat | 86 | 0.454 |
| *sys-1* | β-cat | 153 | 0.555 |
| *wrm-1* | β-cat/APC | 150 | 0.574 |
| *lin-44* | *Wnt* | 117 | 0.489 |
| *egl-20* | *Wnt* | 64 | 0.227 |
| *mom-2* | *Wnt* | 60 | 0.332 |
| *cdc-42* | CDC42 | 49 | 0.420 |

| Gene ID | Protein | Count | I<sub>CG</sub> |
| --- | --- | --- | --- |
| *let-502* | ROCK | 196 | 0.369 |
| *plc-1* | PLC | 230 | 0.446 |
| *plc-2* | PLC | 126 | 0.322 |
| *plc-3* | PLC | 166 | 0.388 |
| *plc-4* | PLC | 71 | 0.168 |
| *tpa-1* | PKC | 505 | 0.274 |
| *pkc-1* | PKC | 85 | 0.118 |
| *pkc-2* | PKC | 86 | 0.084 |
| *pop-1* | APC | 127 | 0.623 |
| *apr-1* | APC | 145 | 0.526 |
| *mom-1* | APC | 46 | 0.106 |
| *mom-5* | APC | 79 | 0.351 |
| *pry-1* | AXIN | 106 | 0.571 |
| *axl-1* | AXIN | 40 | 0.144 |
| *itr-1* | IP3 | 236 | 0.249 |
| *ced-10* | Rac | 76 | 0.601 |
| *mig-2* | Rac | 52 | 0.421 |
| *rac-2* | Rac | 36 | 0.274 |

The function of MELK is related to the regulation of cell size asymmetry and daughter cell fate determination in asymmetric neuroblasts (Cordes et al., 2006). Meanwhile, PKC is involved in signal transduction, with a similar function in *D. melanogaster* (Morrison et al., 2000) and multicellular animals more generally (Plowman et al., 1999). Ecad is involved in cell-cell adhesion and establishing cell polarity (Klompstra et al., 2015). Finally, β-cat is the basis for a novel β-cat/*Wnt* pathway in *C. elegans*, involving both gene loss and duplications that functionally diverge from the canonical Metazoan pathway (Loh et al., 2016). *C. elegans*-specific function involves fewer proteins for both *Wnt* signaling and cell adhesion. This allows for more specific links between gene sequence and protein function, and increased divergence after gene duplication (Jackson and Eisenmann, 2015).

We can also learn something from what does not show up in the group of elevated proteins. For example, Rac is present at intermediate levels in *C. elegans* compared with all protein identities (14), and their candidate genes (*ced-10, mig-2, rac-2*) reveal a moderate to high *I_CG_* value. While Rac is not directly involved in cell differentiation and morphogenesis, it is associated with cytoskeletal modifications and cell migration during axonogenesis (Lundquist, 2001). Proteins such as Calcineurin (Dwivedi et al., 2009), CK-1 (Guillen et al., 2020), and Rho Kinase (Hahman and Schroeter, 2010) has diverse functions in support of cell differentiation such as microtubules and morphogenetic changes (Rho), transcriptional control and *Wnt* signaling (CK-1), and autophagic control (Calcineurin). All these examples involve low to moderate counts for protein identities and moderate *I_CG_* values.

### 4.3 Comparative proteomics

The second part of the proteomic analysis involves a comparison across species with different types of development. Table 4 shows protein counts for four taxa: *C. elegans*, *D. melanogaster*, *M. musculus*, and S. *cerevisiae*. Moving back out to comparisons of protein identities and their counts across species, we can see that the number of proteins in *C. elegans* and *D. melanogaster* (the two examples of mosaic development) exhibit their own unique signatures. One factor driving this difference is variation across pathways and the number of associated genes involved in regulating protein number. For example, *C. elegans* has both a unique β-cat/*Wnt* pathway and no expression of NFAT. Another is the need to maintain a balance of phosphorylation (kinases and phosphatases) in each organism (Plowman et al., 1999; Morrison et al., 2000; Ochoa, 2020).

**Table 4.**
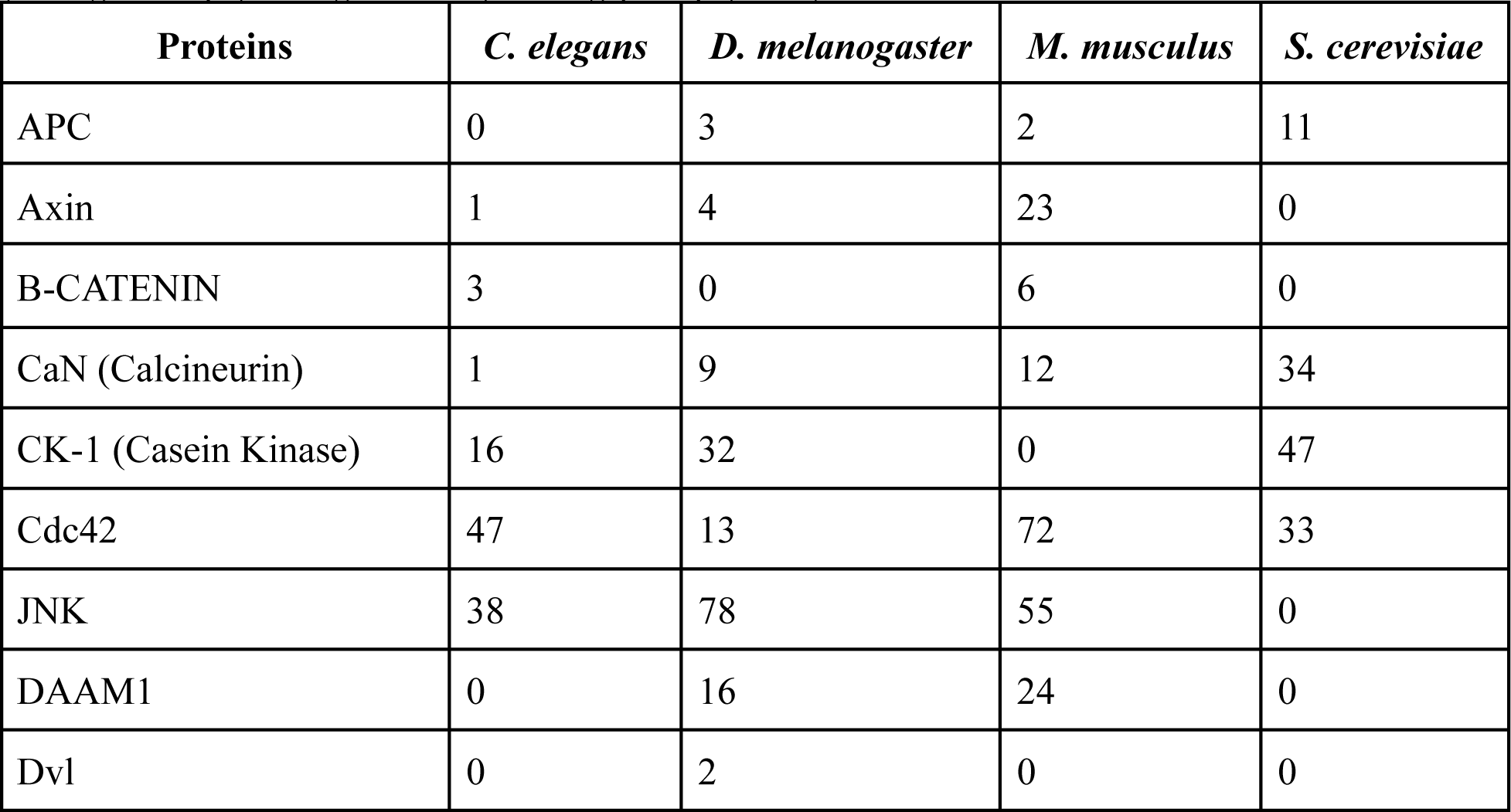
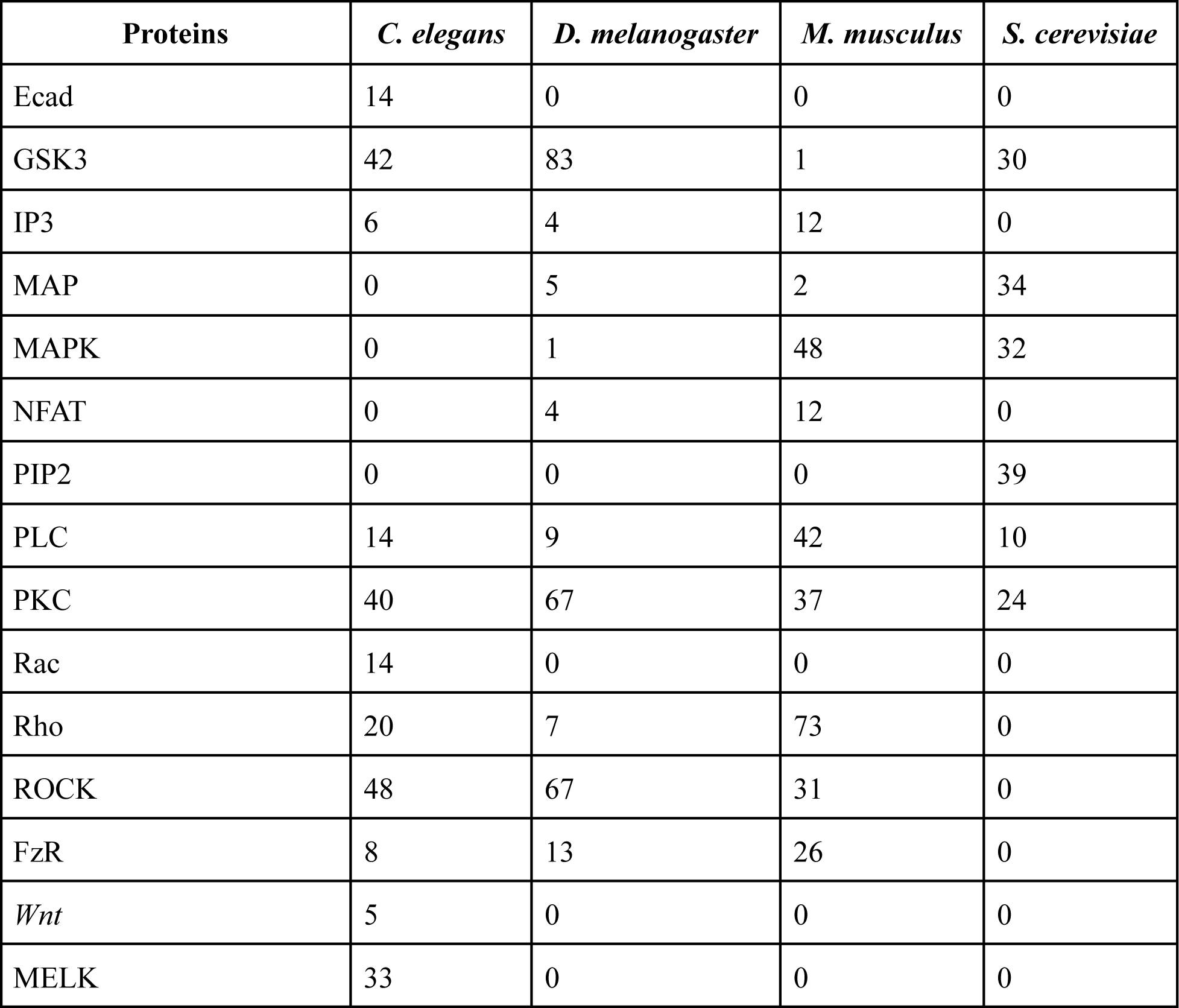
Protein counts for *C. elegans*, *D. melanogaster*, *M. musculus*, and S. *cerevisiae*. Alternate names (paralogs): For *M. musculus*: Fam82a1 (MAP), Mek Kinase 4a (MAPK), NFAT5 (NFAT), Rhodopsin (Rho), ROCK ½ (ROCK), Frizzled (FzR). For *S. cerevisiae*: kar9b (APC), Yck2p (CK-1), Rim11 (GSK3), pkc1p (PKC).

Finally, we can examine protein counts and the differences between mosaic development (*C. elegans and D. melanogaster*) and basic changes in polarity and adhesion (*S. cerevisiae*, see Figures 9A and 9B). Upon normalizing the counts, APC, CaN, CK-1, and MAP/MAPK, and PIP2 are upregulated in S. cerevisiae relative to mosaic development. ROCK and FzR are upregulated in both mosaic species with respect to *S. cerevisiae*. We can also look at differences between mosaic development and an exemplar of regulative development (*M. musculus*) in Table 4 and Figure 9B. A group of six proteins (JNK, PIP2, IP3, FzR, ROCK, and Rho) are present in *C. elegans*, *D. melanogaster*, and *M. Musculus*, but not in *S. cerevisiae.* These six proteins might be considered the most developmentally exclusive. PKC and Cdc42 are present in significant numbers in all species sampled (the most ubiquitous proteins). More common still are proteins present in large numbers in one species, or in all but one species, regardless of developmental status. These include APC (*S. cerevisiae*), Axin (*M. musculus*), CK-1 (*M. musculus*), GSK3 (M. musculus), IP3 (*M. musculus*), MAP (*S. cerevisiae*), IP3 (*M. musculus*), MAP *(S. cerevisiae*), NFAT (*M. musculus*), PIP2 (*S. cerevisiae*), and Rac, *Wnt*, and MELK (*C. elegans*). Some of this (at least in the case of *Wnt*) is an artifact of proteins having different nomenclatures in different species. Some of this also relates to our source of data not being explicitly developmental, but even then yields patterns that follow the different modes of development.

**Figure 9.**
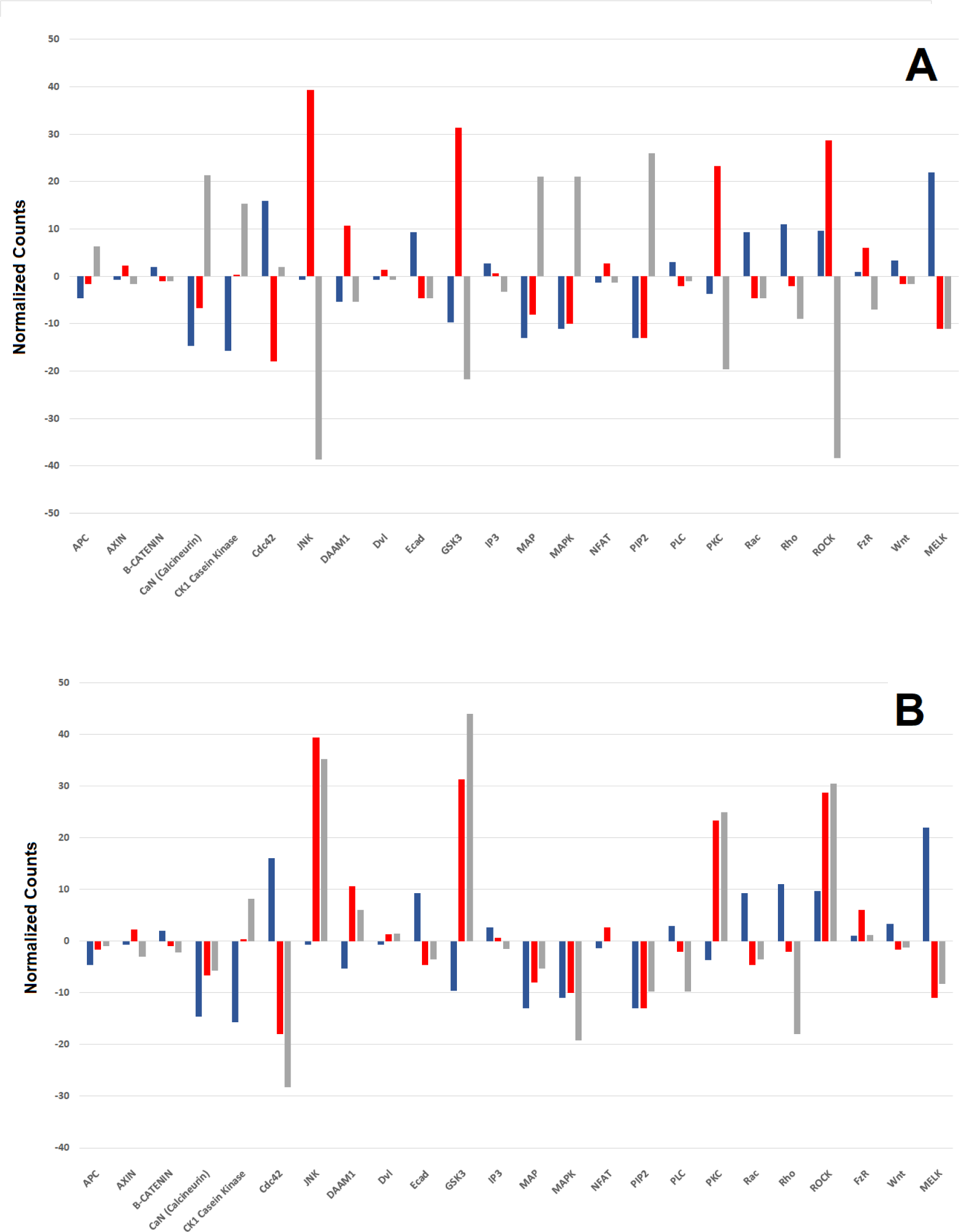
Comparison of mosaic development (*C. elegans* and *D. melanogaster*), regulative development (*M. musculus*), and basic changes in polarity and adhesion (*S. cerevisiae*). A: Normalized counts for *C*. *elegans* (blue), *D. melanogaster* (red), and *S. cerevisiae* (gray). B: Normalized counts for *C. elegans* (blue), *D. melanogaster* (red), and *M. musculus* (gray).

### 4.4 Mapping results to differentiation mechanisms

Now that we have characterized our candidate proteins, we would like to propose a mechanism for differentiation in mosaic development, whether it be asymmetric cell division or terminal differentiation. To do this, we turn to mapping our epigenomic-proteomic line of inquiry onto a computational model. We will use our two MIC models (fully-connected and ordered) to better understand the role of developmental expression for genes associated with our candidate proteins. These modeling results in *C. elegans* will motivate a theoretical discussion on functionally-relevant combinatorial expression across phylogeny.

#### 4.4.1 Fully-connected MIC

The relationship between protein abundance and each fully-connected network (log||A||) is shown in Figure 10. This reveals a slight negative correlation: as protein abundance increases, the determinant of the network decreases. In general, the ||A|| value represents the combination of average RNA copy number and a larger network. Lower average copy number and small networks results in a lower ||A|| value, while a higher average copy number and larger networks result in a higher ||A|| value.

**Figure 10.**
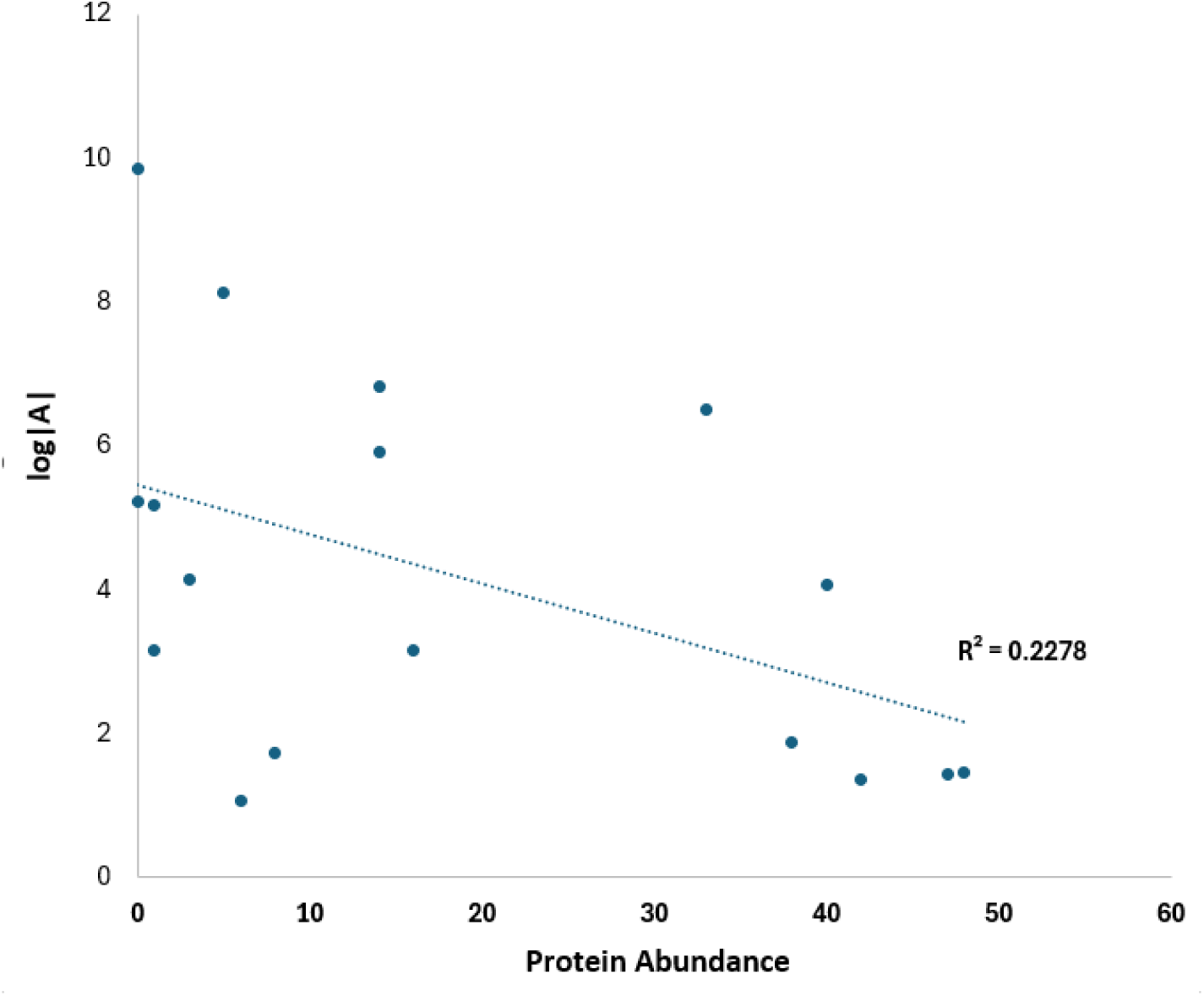
A comparison of protein abundance versus log||A||.

Taking a deeper look at the unweighted fully-connected MICs, we see elevated absolute difference values for a number of genes that contribute to four separate networks. *mom-1* exhibits elevated absolute difference values against all other genes in the APC network. *mom-2* exhibits the same for all other genes in the Wnt network. The same is also true for *axl-1* in the Axin network. Finally, there are elevated absolute value differences between *par-1* and *ham-1* in the MELK network.

#### 4.4.2 Ordered MIC

For each protein’s ordered MIC, we take each four compartment element and evaluate the increase between the early- and mid-developmental condition. An increase between early- and mid-development is representative of an expansion wave signal, while a decrease between early- and mid-development is representative of a contraction wave signal. The ordered network is characterized by chromosomal and genomic order, their corresponding epigenetic weights, and the direction of average gene expression difference between early- and mid-development. The results for 17 networks are shown in Figure 11 (diagrammatic) and Appendix B (numeric values).

**Figure 11.**
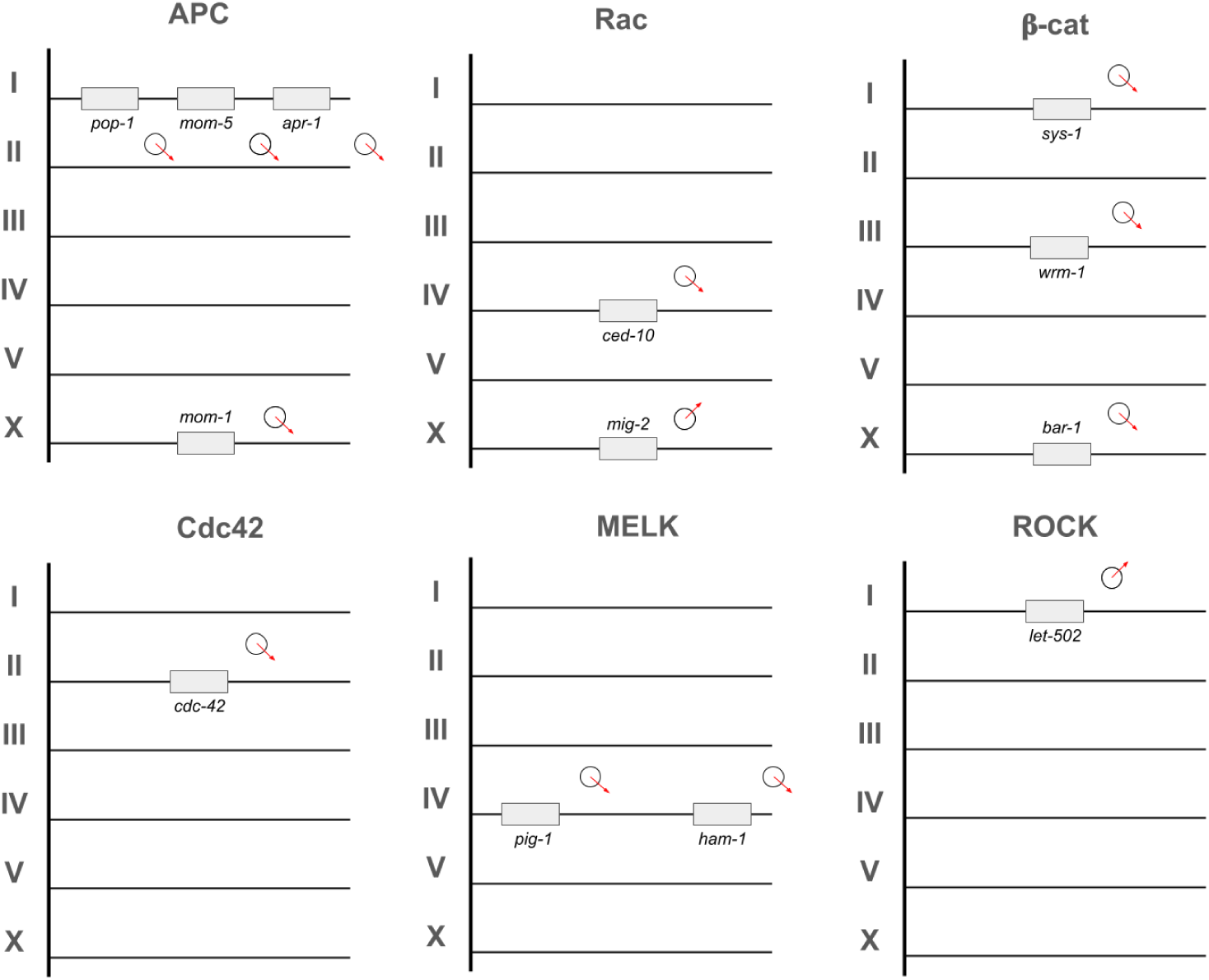
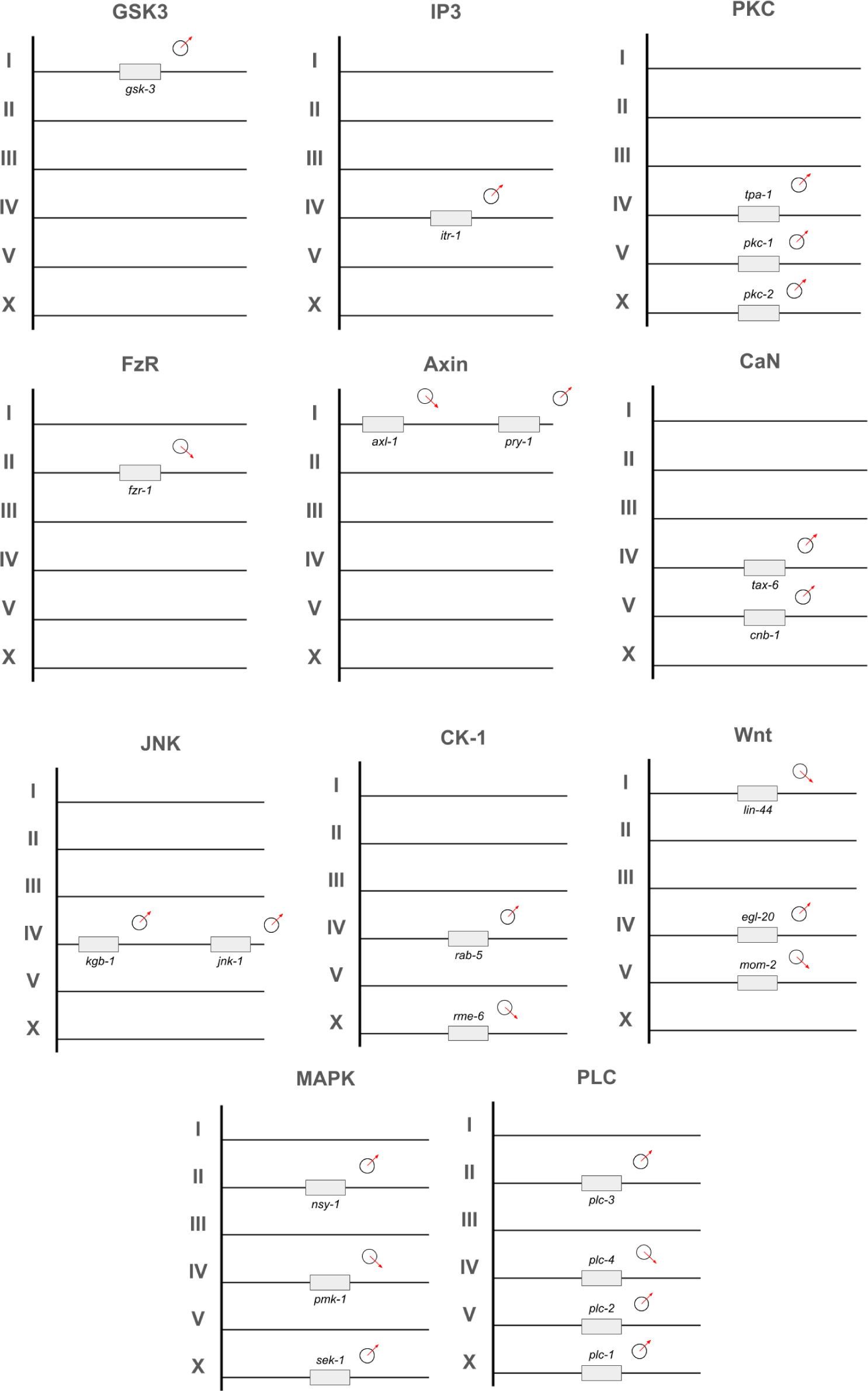
The 17 ordered MICs, the chromosomal location of each gene, and the direction of change in average gene expression across embryonic development. Upward arrows indicate an expansion wave signal, and downward arrows indicate contraction wave signals.

A map of the accumulated signals for each network is shown in Figure 11. While the signals for genes on the same chromosome are equally represented by expansion and contraction waves (I = 4 expansion, 5 contraction; II = 2 expansion, 2 contraction; III = 0 expansion, 1 contraction; IV = 7 expansion, 5 contraction; V = 3 expansion, 1 contraction; and X = 4 expansion, 3 contraction). Yet for genes associated with individual proteins, we see asymmetric signals dominated by expansion (PKC = 3 expansion, 0 contraction, abundance of 40); CaN and JNK = 2 expansion, JNK abundance of 38; and PLC = 3 expansion, 1 contraction, abundance of 14) and contraction (APC = 4 contraction, abundance of 0; Rac = 3 contraction, abundance of 14; and MELK = 2 contraction, abundance of 33) signals. While not all genetic contributions to each protein have been accounted for, these genes represent the most characterized contributions to the protein, and perhaps even predicting the role of each gene in differentiation waves.

Table 5 shows the ensemble values yielded by the ordered MICs for each of our 17 proteins. Each gene provides a facilitative (weighted upregulation between early- and mid-development) or suppressive (weighted downregulation between early- and mid-development) signal. For example, the MAPK contains three genes with an ensemble that is primarily (0.66) facilitative: two genes contributing to that facilitative signal (*sek-1*, *nsy-1*) and one that is suppressive (*pmk-1*). This yields ensembles for both proteins (within a specific ordered MIC) and for chromosomes across proteins (between ordered MICs). Most protein-specific ordered MICs yield either unified facilitative or suppressive. In fact, only Rac, CK-1, Axin, Wnt, and MAPK yield a heterogeneous ensemble (Wnt being biased towards suppression, MAPK being biased towards facilitation). Ordered MIC agnostic chromosomal ensembles also yield heterogeneous results: Chromosomes I, III, and X being biased towards suppression, and Chromosomes II, IV, and V biased towards facilitation. A combination of within- and between ordered MIC signal processing may more fully describe the outcomes observed in developmental phenotypes.

**Table 5.** Ensemble values for each protein and chromosome. An ensemble is the relative
contribution of each gene as a facilitative or suppressive signal.

| Protein/Chromosome ID | Facilitative | Suppressive |
| --- | --- | --- |
| APC | 0.0 | 1.0 |
| Rac | 0.5 | 0.5 |
| β-cat | 1.0 | 0.0 |
| Cdc42 | 0.0 | 1.0 |
| MELK | 0.0 | 1.0 |
| ROCK | 1.0 | 0.0 |
| GSK3 | 1.0 | 0.0 |
| IP3 | 1.0 | 0.0 |
| PKC | 1.0 | 0.0 |
| FzR | 0.0 | 1.0 |
| Axin | 0.5 | 0.5 |
| CaN | 1.0 | 0.0 |
| JNK | 1.0 | 0.0 |
| CK-1 | 0.5 | 0.5 |
| Wnt | 0.33 | 0.67 |
| MAPK | 0.66 | 0.33 |
| I | 0.33 | 0.67 |
| II | 0.67 | 0.33 |
| III | 0.0 | 1.0 |
| IV | 0.58 | 0.42 |
| V | 0.75 | 0.25 |
| X | 0.43 | 0.57 |

### 4.5 Combinatorial aspects of differentiation tree topology

There is also a connection between the combinatorial aspects of the regulatory computational models and the topology of a differentiation tree. As the differentiation code predicts protein regulation and localization of structure, the copy, paste, and delete operations imposed by evolutionary change are reflected in the differentiation tree. Our simplest representation of this differentiation tree to regulatory computational model mapping is shown in Figures 12A-12D. Recall that in mosaic development, differentiation trees are simply reorganized lineage trees. In regulative development, the corresponding changes in tissue differentiation are decoupled from the lineage tree.

**Figure 12.**
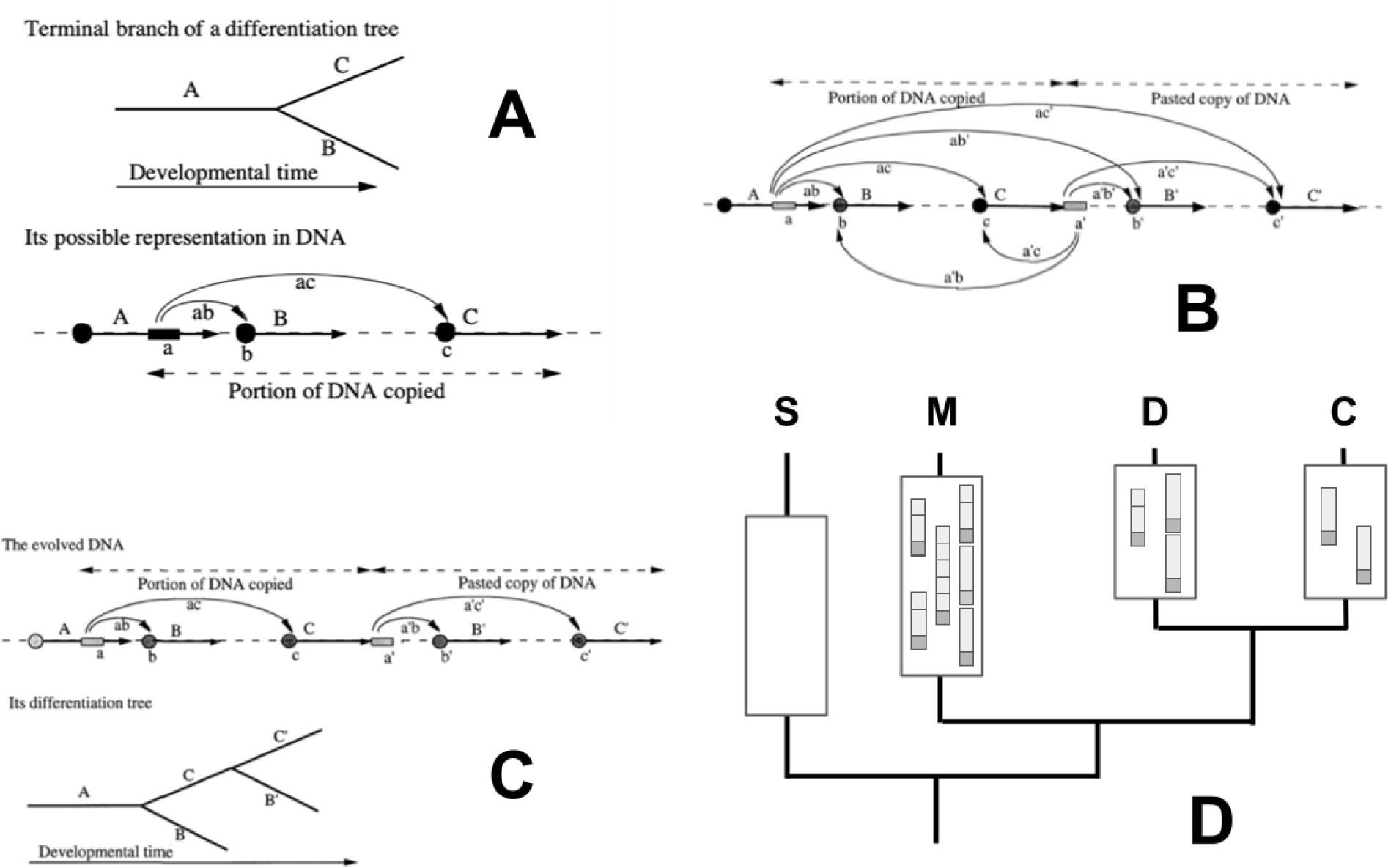
Mapping genomic operations to the differentiation tree, and how the resulting gene maps to hypothetical regulatory computational model evolution on a phylogeny for our model organisms. A) model for how a gene duplication (a single promoter region regulating two coding regions, one copied from the other) can serve as the basis for a differentiation tree branch. Taken from Figure 24 in (Gordon, 1999). Reproduced with permission of World Scientific Press [to be requested]. B) State of the DNA after duplication. Taken from Figure 25 in (Gordon, 1999). Permission of World Scientific Press [to be requested]. C) Copy and paste operation that leads to two levels of a differentiation tree Taken from Figure 25 in (Gordon, 1999). Permission of World Scientific Press [to be requested]. D) a phylogeny showing *S. cerevisiae* [S], *M. musculus* [M], *D. melanogaster* [D], and *C. elegans* [C] as per their phylogenetic position. Insets on each terminal branch show changes in the same regulatory computational model for the corresponding organism.

In Figure 12A, we can see a simple representation of a gene with promoter (a) and coding regions (b and c). As in Figure 3, this produces several alternate transcripts (ab and ac). In the corresponding differentiation tree diagram, tissue type A erects cell state splitters in all its cells that resolve their mechanical instabilities in all its cells as either cell type B or C. The cells resulting from B and C are considered terminally differentiated cell types in this example. The activator for gene cascade A, a, sends out one of two signals in each cell: either ab or ac, depending on whether the cell has participated in an expansion or a contraction wave (Bjorklund and Gordon, 1993; Bjorklund and Gordon, 1994). These signals activate either gene cascade B or C, respectively (Gordon, 1999).

In Figure 12B, we assume the simplest case, namely that the copied terminal branch is inserted at the end of the formerly terminal C cascade (a reduplication *sensu* Darlington, 1958), extending the terminal branch from which it was copied, in such a way that activation of cascade C also leads to the copy of the activator a, namely a’, also being activated. This in turn activates either B’ or C’ through signals a’b’ or a’c’, respectively. However, immediately after such a duplication, crosstalk cannot be avoided. Thus, the activator a will have a direct link with initiators b’ and c’, through signals ab’ and ac’. Similarly, signal a’ will also react in a backward fashion to cascades B and and C through the signals a’b and and a’c, respectively. In terms of molecular structure, all combinations are equivalent (Gordon, 1999).

Figure 12C shows a copy and paste operation as the basis for coevolution after a duplication event. Activator **a’** is assumed to coevolve with initiators **b’** and **c’**, so that a new, cleanly separated terminal branch of the differentiation tree results. In this diagram, the old cascades are shown unchanged, though they, too, could continue evolving. Thus, we can end up with a’ ≠ a, b’ ≠ c, c’ ≠ c, a’b’ ≠ ab, a’c’ ≠ ac, B’ ≠ B, and C’ ≠ C (Gordon, 1999). In Figure 12D, we provide an example of how a single regulatory computational model evolves not within the context of a single species differentiation tree, but across our four model species. This gives us a set of phylogenetic relationships that also show how components of the regulatory computational model are lost, duplicated, copied, and otherwise moved around the genome for each of our model species. This example is not based on the substrate for any one protein count but shows more generally how we expect that our regulatory computational model will produce homologs and paralogs, which were reported for all our protein comparisons.

## 5 Discussion

In this paper, we introduce the reader to the problem of differentiation in embryological development. We focus on the nematode *C. elegans* due to its wealth of data and amenable biological properties. The differentiation code is derived from the differentiation tree and together provide a theoretical basis for understanding how cell differentiation translates into a phenotype. Figure 1 shows how differentiation waves can be mapped to two anatomical axes of the embryo, providing a route to understanding differentiation waves using techniques such as spatial genomics. To understand the molecular counterparts of differentiation according to our hypothesis, we focus on gene expression, epigenomics, and proteomics in *C. elegans* while taking a comparative approach to widen the scope of our proteomic account. In the comparative analysis, proteins from four organisms are used to represent mosaic development, regulative development, and basic changes in polarity and adhesion. Our analysis of protein expression suggests patterns of protein abundance and absence that are dependent on species and type of development. To bring together gene expression, epigenetic, and proteomic data for a host of candidate genes associated with our candidate proteins, we build two different types of MIC using data from *C. elegans*. The fully-connected MIC captures differences in the potential interactions of genes, while the ordered MIC captures the linear contributions of each gene to a potential developmental outcome.

There are several caveats to our empirical analyses. One is that the connection between our candidate genes and protein list are limited to those identified in the literature. There may be other genes involved in the corresponding proteins not included in the epigenome analysis. However, these genes have the largest effect, and should provide the best snapshot of the proposed relationships. The second is that several of our gene sequences are contigs rather than a full sequence. Since we do not distinguish between promoter regions and exons, this is not a crucial detail. Interestingly, in a few cases, there are coding regions of other genes nested in at least one of our genes of interest (*rac-2* being nested within the length of *ced-10*). As we are working from a well-annotated reference genome, this type of complexity might direct future refinement of our model. Additional future work may be done by enhancing our knowledge of these MICs and protein abundances with additional bioinformatic analysis from publicly-available databases. Future work might also consider alternate approaches to MICs that describe the link between gene expression, proteins, and tissue differentiation. One emerging method called interaction mapping allows us to explore chemical-genetic interactions (Braberg et.al, 2022). A means to explore the intersection of network and structural biology would enable more precise information about gene networks and proteins explicitly involved in triggering differentiation waves. Another model that might be fruitful are multi-objective process models and the more general hybrid cybernetic models (Aboulmouna et.al, 2020), where regulatory processes are recast as a set of optimization objectives. Hybrid cybernetic models in particular serve as alternatives to boolean GRNs, and serve to constrain the activity of various biochemical networks in biologically-plausible ways (Young et.al, 2008; Song and Ramkrishna, 2010).

Other mechanisms for controlling cell proliferation can also provide insights into which MICs provide signals for differentiation waves. Spatial constraints coupled with mechanical feedback from the local cell population can control the rate of cell proliferation (Streichan et.al, 2014), leading to the formation and growth of tissues (see Figure 13C). In limited cases, we can look at the effects of genetic drift on proteins that are the products of gene duplication. By looking more deeply at the Blastp results, we found that FzR (Frizzled) is subject to drift after duplication. FzR is an interesting example (Figure 13B), as it is intensely involved in developmental processes such as epidermal cell polarity, formation of synapses, and the regulation of proliferation (Huang and Klein, 2004). As with most types of developmental proteins, we expect FzR to be conserved. But protein counts and the *I_CG_* values for *fzr-1* (*frizzled* gene in *C. elegans*) do not stand out in our sample.

**Figure 13.**
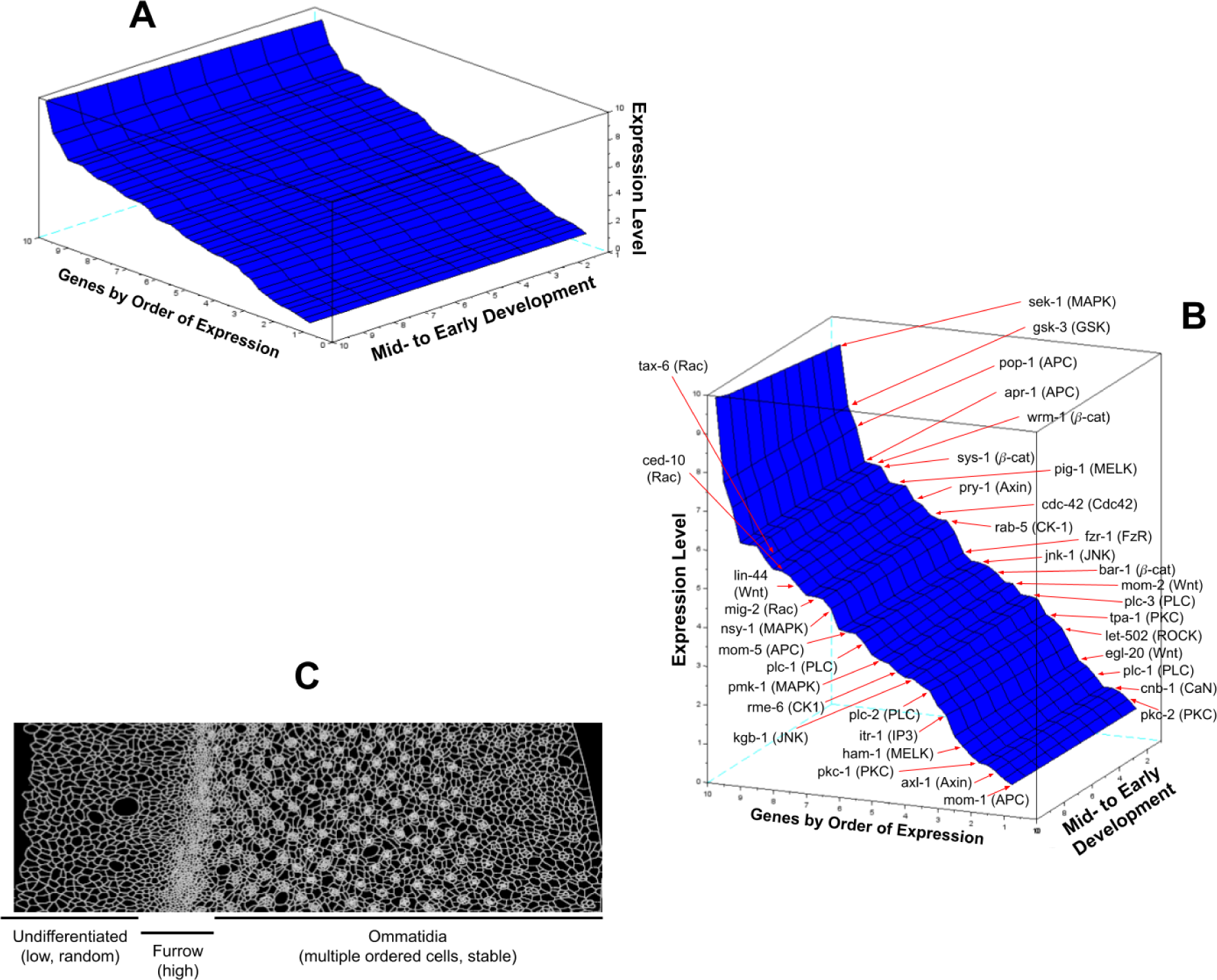
A) A 3-D surface plot showing individual genes sorted by lowest to highest expression level and changes in expression between early and mid-development. B) An alternate angular view of this same 3-D surface plot labeled with the identity of individual genes. C) expected phenotypic consequences with respect to fully-connected and ordered MICs as shown for a developing *Drosophila* compound eye (imaginal disc cells differentiating into ommatidia). Description in parentheses is the phenotypic structure and expected mode of gene expression for a certain region of tissue. Values along the mid-to early development axis are interpolated using a smoothing technique (midpoints between the early- and mid-development measurements as reported). Image of imaginal disc is from Alicea et.al (2018).

The top to bottom undulations in Figure 13 (outputs per gene axis) demonstrate the stepwise increase in expression as ordered by early development gene expression values. This yields large intervals towards the top of the graph, typical of more highly expressed genes. The lateral undulations (mid-to early development axis) results from variation in expression for single genes over time. We observed a variety of differentiation signals for Wnt, MAPK, Rac, Axin, and CK-1, which may be suggestive for tissue-wide roles. Purely facilitative signals characterize *β*-cat, ROCK, GSK3, IP3, PKC, CaN, and JNK expression, while purely suppressive signals characterize APC, Cdc42, MELK, and FzR. Figure 13C allows us to consider how this might map to an active differentiation process according to the wave mechanism. For undifferentiated cells (left-hand side of the imaginal disc), low/random gene expression of our target genes requires suppressive signals to dominate. The furrow (mid-section of imaginal disk) is characterized by high expression, and requires a predominance of facilitative signals. As a collection of different cell types, differentiated ommatidia structures (right-hand side of the imaginal disk) require a heterogeneity of signals,

As we have given only a quick sketch of regulatory complexity in our computational models, we have still not made specific predictions about protein activity. Future work will require a broader range of experimental data and a better representation of all genes involved in the regulatory computational model that underlie the predicted phenotype is also needed. Here we propose that this could be done by examining the genomes of fully DNA sequenced organisms for which we also know the differentiation tree. While we have a differentiation tree for the beginning of embryogenesis for the axolotl, *Ambystoma mexicanum* (Figure 2), and its huge genome has been sequenced (Keinath et al., 2015), we do not know the full differentiation tree topology. However, this paper provides us with a way to derive and use differentiation codes for a wide variety of contexts.

## Acknowledgements

We thank Andrei Igamberdiev for his careful comments, and the DevoWorm group for their feedback.

## Appendix

**Appendix A.** Matrices for 17 fully-connected MICs and their associated weight matrices.

| Fully-connected MIC |  |  |  |  | MIC Weights |  |  |  |  |
| --- | --- | --- | --- | --- | --- | --- | --- | --- | --- |
| **APC** | *pop-1* | *apr_1* | *mom-1* | *mom-5* | **APC** | *pop-1* | *apr_1* | *mom-1* | *mom-5* |
| *pop-1* | 64.8 | 22 | 1264.8 | 12 | *pop-1* | 0.53 | 0.33 | 0.18 | 0.06 |
| *apr_1* | 22 | 42.8 | 1218.7 | 10 | *apr_1* | 0.33 | 0.62 | 0.22 | 0.07 |
| *mom-1* | 1264.8 | 1318.7 | 1361.5 | 1308.7 | *mom-1* | 0.19 | 0.22 | 0.35 | 0.04 |
| *mom-5* | 12 | 10 | 1308.7 | 52.8 | *mom-5* | 0.06 | 0.07 | 0.04 | 0.11 |
| **Rac** | *ced-10* | *mig-2* | *rac-2* |  | **Rac** | *ced-10* | *mig-2* | *rac-2* |  |
| *ced-10* | 72.7 | 65.3 | 72.7 |  | *ced-10* | 0.60 | 0.25 | 0.16 |  |
| *mig-2* | 65.3 | 138 | 138 |  | *mig-2* | 0.25 | 0.42 | 0.12 |  |
| *rac-2* | 72.7 | 138 | 0 |  | *rac-2* | 0.16 | 0.12 | 0.27 |  |
| **β-cat** | *bar-1* | *sys-1* | *wrm-1* |  | **β-cat** | *bar-1* | *sys-1* | *wrm-1* |  |
| *bar-1* | 63.2 | 99.7 | 28.7 |  | *bar-1* | 0.45 | 0.25 | 0.26 |  |
| *sys-1* | 99.7 | 162.9 | 71 |  | *sys-1* | 0.25 | 0.56 | 0.32 |  |
| *wrm-1* | 28.7 | 71 | 91.9 |  | *wrm-1* | 0.26 | 0.32 | 0.57 |  |
| **Cdc42** | *cdc-42* |  |  |  | **Cdc42** | *cdc-42* |  |  |  |
| *cdc-42* | 27.8 |  |  |  | *cdc-42* | 0.42 |  |  |  |
| **MELK** | *pig-1* | *par-1* | *ham-1* |  | **MELK** | *pig-1* | *par-1* | *ham-1* |  |
| *pig-1* | 117.6 | 109.9 | 91.3 |  | *pig-1* | 0.58 | 0.14 | 0.08 |  |
| *par-1* | 109.9 | 7.7 | 201.2 |  | *par-1* | 0.14 | 0.25 | 0.03 |  |
| *ham-1* | 91.3 | 201.2 | 208.9 |  | *ham-1* | 0.08 | 0.03 | 0.14 |  |
| **ROCK** | *let-502* |  |  |  | **ROCK** | *let-502* |  |  |  |
| *let-502* | 29.8 |  |  |  | *let-502* | 0.369 |  |  |  |
| **GSK3** | *gsk-3* |  |  |  | **GSK3** | *gsk-3* |  |  |  |
| *gsk-3* | 22.6 |  |  |  | *gsk-3* | 0.596 |  |  |  |
| **IP3** | *itr-1* |  |  |  | **IP3** | *itr-1* |  |  |  |
| *itr-1* | 11.4 |  |  |  | *itr-1* | 0.249 |  |  |  |
| **PKC** | *tpa-1* | *pkc-1* | *pkc-2* |  | **PKC** | *tpa-1* | *pkc-1* | *pkc-2* |  |
| *tpa-1* | 31.8 | 11.9 | 4.6 |  | *tpa-1* | 0.27 | 0.03 | 0.02 |  |
| *pkc-1* | 11.9 | 19.9 | 7.3 |  | *pkc-1* | 0.03 | 0.12 | 0.01 |  |
| *pkc-2* | 4.6 | 7.3 | 27.2 |  | *pkc-2* | 0.02 | 0.01 | 0.08 |  |
| **FzR** | *fzr-1* |  |  |  | **FzR** | *fzr-1* |  |  |  |
| *fzr-1* | 55.6 |  |  |  | *fzr-1* | 0.378 |  |  |  |
| **Axin** | *pry-1* | *axl-1* |  |  | **Axin** | *pry-1* | *axl-1* |  |  |
| *pry-1* | 58.3 | 416.9 |  |  | *pry-1* | 0.57 | 0.08 |  |  |
| *axl-1* | 416.9 | 475.2 |  |  | *axl-1* | 0.08 | 0.14 |  |  |
| **CaN** | *tax-6* | *cnb-1* |  |  | **CaN** | *tax-6* | *cnb-1* |  |  |
| *tax-6* | 11.1 | 45.8 |  |  | *tax-6* | 0.52 | 0.06 |  |  |
| *cnb-1* | 45.8 | 56.9 |  |  | *cnb-1* | 0.06 | 0.12 |  |  |
| **JNK** | *jnk-1* | *kgb-1* |  |  | **JNK** | *jnk-1* | *kgb-1* |  |  |
| *jnk-1* | 105.8 | 65.7 |  |  | *jnk-1* | 0.46 | 0.18 |  |  |
| *kgb-1* | 65.7 | 40.1 |  |  | *kgb-1* | 0.18 | 0.39 |  |  |
| **CK-1** | *rme-6* | *rab-5* |  |  | **CK-1** | *rme-6* | *rab-5* |  |  |
| *rme-6* | 70 | 33.3 |  |  | *rme-6* | 0.40 | 0.16 |  |  |
| *rab-5* | 33.3 | 36.7 |  |  | *rab-5* | 0.16 | 0.40 |  |  |
| **Wnt** | *lin-44* | *egl-20* | *mom-2* |  | **Wnt** | *lin-44* | *egl-20* | *mom-2* |  |
| *lin-44* | 387 | 260.3 | 1357.3 |  | *lin-44* | 0.49 | 0.11 | 0.16 |  |
| *egl-20* | 260.3 | 126.7 | 1617.6 |  | *egl-20* | 0.11 | 0.23 | 0.08 |  |
| *mom-2* | 1357.3 | 1617.6 | 1744.3 |  | *mom-2* | 0.16 | 0.08 | 0.33 |  |
| **MAPK** | *pmk-1* | *nsy-1* | *sek-1* |  | **MAPK** | *pmk-1* | *nsy-1* | *sek-1* |  |
| *pmk-1* | 78.7 | 59.1 | 18.2 |  | *pmk-1* | 0.31 | 0.13 | 0.28 |  |
| *nsy-1* | 59.1 | 19.6 | 40.9 |  | *nsy-1* | 0.13 | 0.43 | 0.38 |  |
| *sek-1* | 18.2 | 40.9 | 60.5 |  | *sek-1* | 0.28 | 0.38 | 0.89 |  |
| **PLC** | *plc-1* | *plc-2* | *plc-3* | *plc-4* | **PLC** | *plc-1* | *plc-2* | *plc-3* | *plc-4* |
| *plc-1* | 28.9 | 75.8 | 27.9 | 5.4 | *plc-1* | 0.45 | 0.14 | 0.17 | 0.07 |
| *plc-2* | 75.8 | 104.7 | 47.9 | 70.4 | *plc-2* | 0.14 | 0.32 | 0.12 | 0.05 |
| *plc-3* | 27.9 | 47.9 | 56.8 | 22.5 | *plc-3* | 0.17 | 0.12 | 0.39 | 0.07 |
| *plc-4* | 5.4 | 70.4 | 22.5 | 34.3 | *plc-4* | 0.07 | 0.05 | 0.07 | 0.17 |

**Appendix B.**
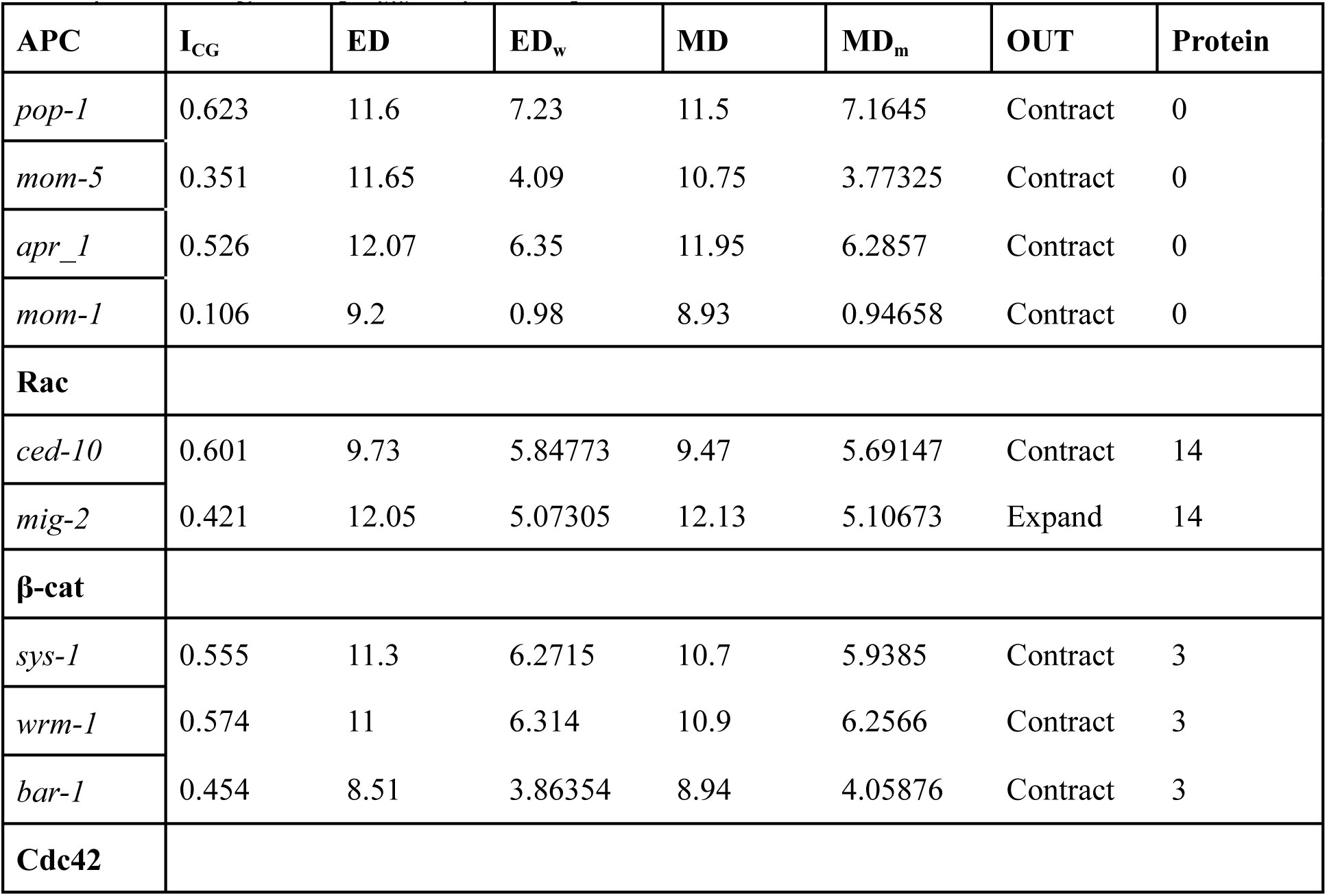
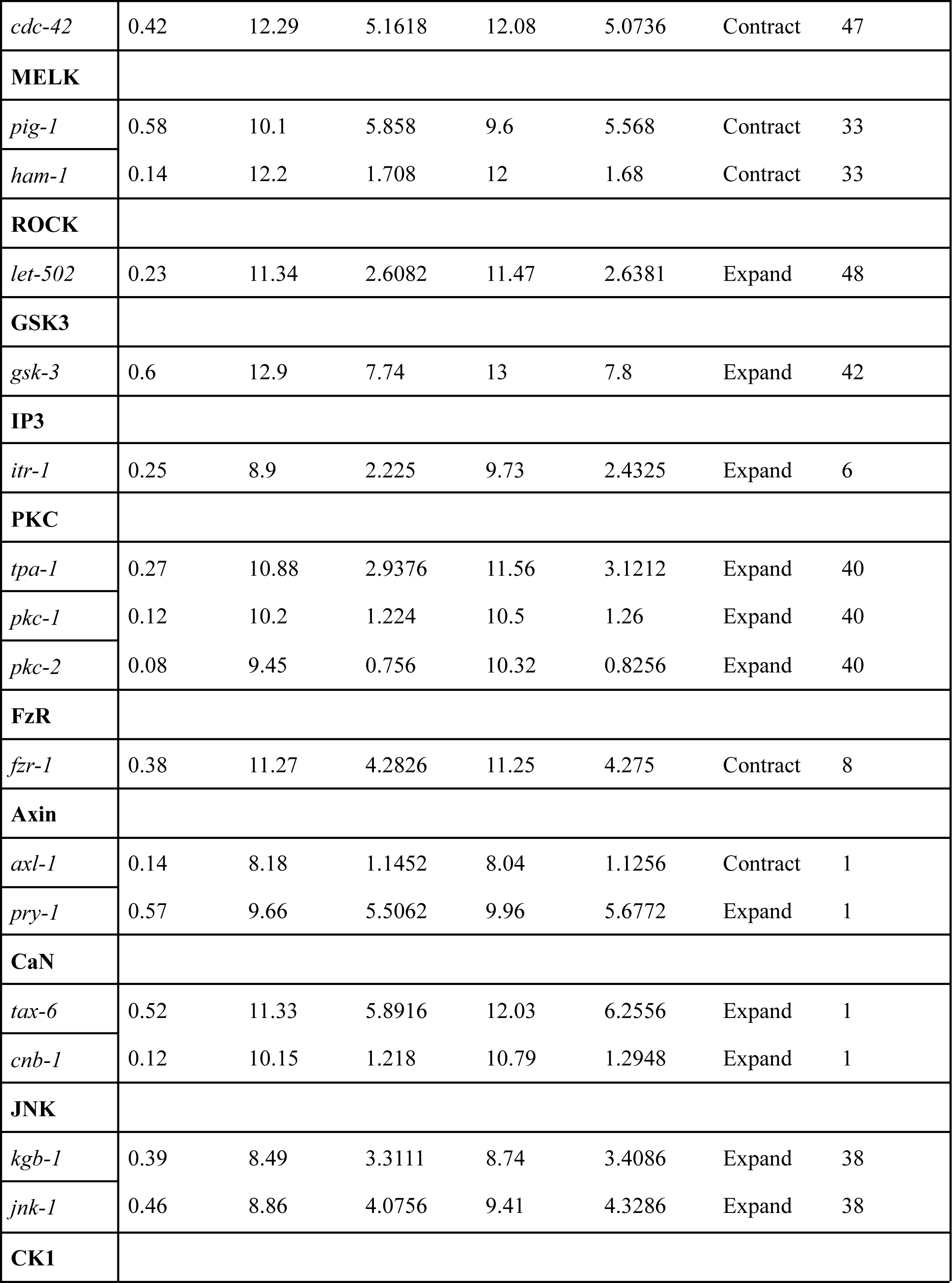
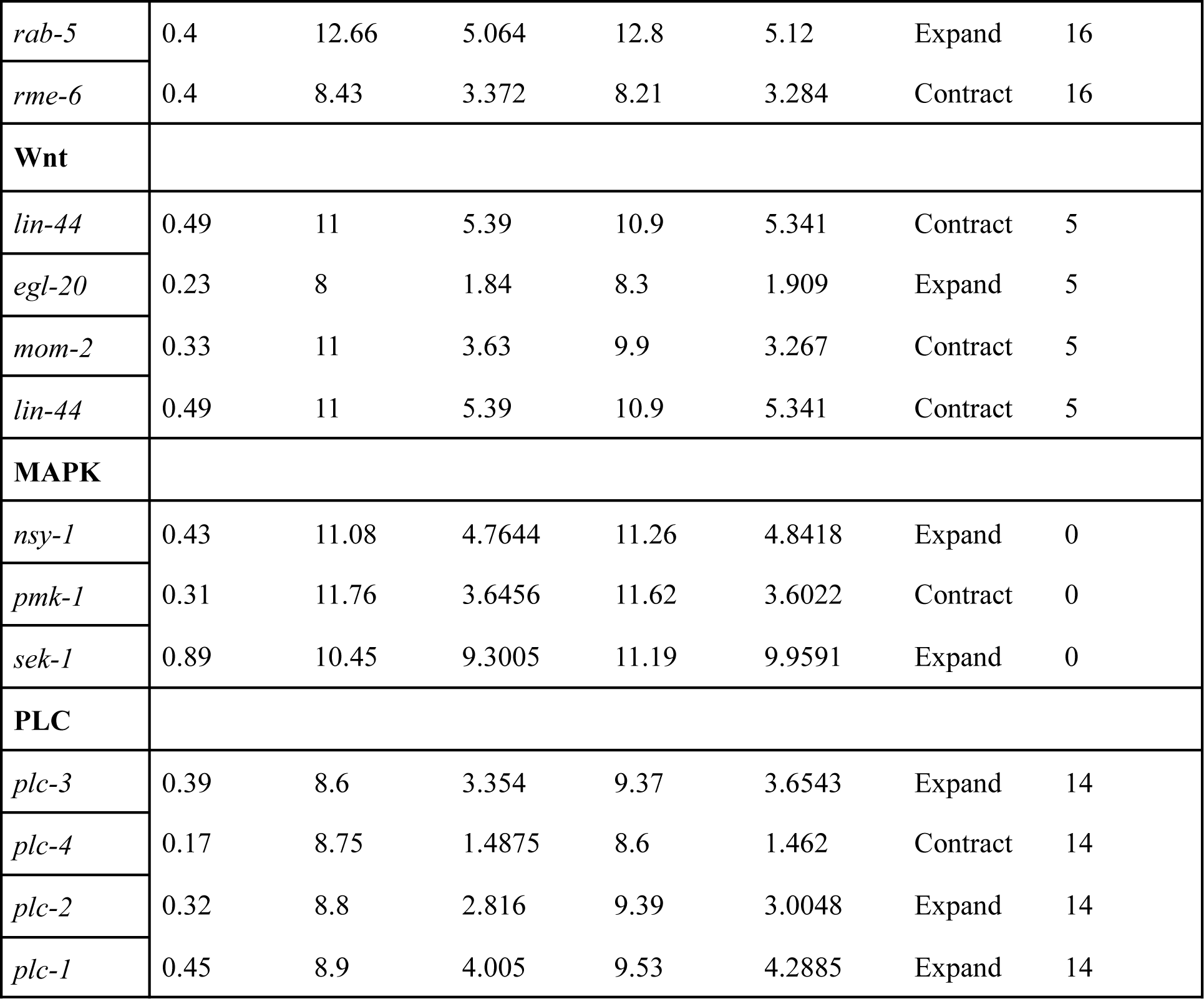
Matrices for 17 ordered MICs with associated protein abundance counts. OUT represents the output of each element as an expansion wave signal (Expand) or contraction wave signal (Contract), while ED_w_ and MD_w_ represent early development weighted by I_CG_ and early development weighted by I_CG_, respectively.

| **APC** | **I<sub>CG</sub>** | **ED** | **ED<sub>w</sub>** | **MD** | **MD<sub>m</sub>** | **OUT** | **Protein** |
| --- | --- | --- | --- | --- | --- | --- | --- |
| *pop-1* | 0.623 | 11.6 | 7.23 | 11.5 | 7.1645 | Contract | 0 |
| *mom-5* | 0.351 | 11.65 | 4.09 | 10.75 | 3.77325 | Contract | 0 |
| *apr_1* | 0.526 | 12.07 | 6.35 | 11.95 | 6.2857 | Contract | 0 |
| *mom-1* | 0.106 | 9.2 | 0.98 | 8.93 | 0.94658 | Contract | 0 |
| **Rac** |  |  |  |  |  |  |  |
| *ced-10* | 0.601 | 9.73 | 5.84773 | 9.47 | 5.69147 | Contract | 14 |
| *mig-2* | 0.421 | 12.05 | 5.07305 | 12.13 | 5.10673 | Expand | 14 |
| **β-cat** |  |  |  |  |  |  |  |
| *sys-1* | 0.555 | 11.3 | 6.2715 | 10.7 | 5.9385 | Contract | 3 |
| *wrm-1* | 0.574 | 11 | 6.314 | 10.9 | 6.2566 | Contract | 3 |
| *bar-1* | 0.454 | 8.51 | 3.86354 | 8.94 | 4.05876 | Contract | 3 |
| **Cdc42** |  |  |  |  |  |  |  |
| *cdc-42* | 0.42 | 12.29 | 5.1618 | 12.08 | 5.0736 | Contract | 47 |
| **MELK** |  |  |  |  |  |  |  |
| *pig-1* | 0.58 | 10.1 | 5.858 | 9.6 | 5.568 | Contract | 33 |
| *ham-1* | 0.14 | 12.2 | 1.708 | 12 | 1.68 | Contract | 33 |
| **ROCK** |  |  |  |  |  |  |  |
| *let-502* | 0.23 | 11.34 | 2.6082 | 11.47 | 2.6381 | Expand | 48 |
| **GSK3** |  |  |  |  |  |  |  |
| *gsk-3* | 0.6 | 12.9 | 7.74 | 13 | 7.8 | Expand | 42 |
| **IP3** |  |  |  |  |  |  |  |
| *itr-1* | 0.25 | 8.9 | 2.225 | 9.73 | 2.4325 | Expand | 6 |
| **PKC** |  |  |  |  |  |  |  |
| *tpa-1* | 0.27 | 10.88 | 2.9376 | 11.56 | 3.1212 | Expand | 40 |
| *pkc-1* | 0.12 | 10.2 | 1.224 | 10.5 | 1.26 | Expand | 40 |
| *pkc-2* | 0.08 | 9.45 | 0.756 | 10.32 | 0.8256 | Expand | 40 |
| **FzR** |  |  |  |  |  |  |  |
| *fzr-1* | 0.38 | 11.27 | 4.2826 | 11.25 | 4.275 | Contract | 8 |
| **Axin** |  |  |  |  |  |  |  |
| *axl-1* | 0.14 | 8.18 | 1.1452 | 8.04 | 1.1256 | Contract | 1 |
| *pry-1* | 0.57 | 9.66 | 5.5062 | 9.96 | 5.6772 | Expand | 1 |
| **CaN** |  |  |  |  |  |  |  |
| *tax-6* | 0.52 | 11.33 | 5.8916 | 12.03 | 6.2556 | Expand | 1 |
| *cnb-1* | 0.12 | 10.15 | 1.218 | 10.79 | 1.2948 | Expand | 1 |
| **JNK** |  |  |  |  |  |  |  |
| *kgb-1* | 0.39 | 8.49 | 3.3111 | 8.74 | 3.4086 | Expand | 38 |
| *jnk-1* | 0.46 | 8.86 | 4.0756 | 9.41 | 4.3286 | Expand | 38 |
| **CK1** |  |  |  |  |  |  |  |
| *rab-5* | 0.4 | 12.66 | 5.064 | 12.8 | 5.12 | Expand | 16 |
| *rme-6* | 0.4 | 8.43 | 3.372 | 8.21 | 3.284 | Contract | 16 |
| **Wnt** |  |  |  |  |  |  |  |
| *lin-44* | 0.49 | 11 | 5.39 | 10.9 | 5.341 | Contract | 5 |
| *egl-20* | 0.23 | 8 | 1.84 | 8.3 | 1.909 | Expand | 5 |
| *mom-2* | 0.33 | 11 | 3.63 | 9.9 | 3.267 | Contract | 5 |
| *lin-44* | 0.49 | 11 | 5.39 | 10.9 | 5.341 | Contract | 5 |
| **MAPK** |  |  |  |  |  |  |  |
| *nsy-1* | 0.43 | 11.08 | 4.7644 | 11.26 | 4.8418 | Expand | 0 |
| *pmk-1* | 0.31 | 11.76 | 3.6456 | 11.62 | 3.6022 | Contract | 0 |
| *sek-1* | 0.89 | 10.45 | 9.3005 | 11.19 | 9.9591 | Expand | 0 |
| **PLC** |  |  |  |  |  |  |  |
| *plc-3* | 0.39 | 8.6 | 3.354 | 9.37 | 3.6543 | Expand | 14 |
| *plc-4* | 0.17 | 8.75 | 1.4875 | 8.6 | 1.462 | Contract | 14 |
| *plc-2* | 0.32 | 8.8 | 2.816 | 9.39 | 3.0048 | Expand | 14 |
| *plc-1* | 0.45 | 8.9 | 4.005 | 9.53 | 4.2885 | Expand | 14 |

